# Ice formation and its elimination in cryopreservation of bovine oocytes

**DOI:** 10.1101/2023.11.15.567270

**Authors:** Abdallah W. Abdelhady, David W. Mittan-Moreau, Patrick L. Crane, Matthew J. McLeod, Soon Hon Cheong, Robert E. Thorne

**Author notes:** **Corresponding authors:** Robert E. Thorne, Soon Hon Cheong **Email:**. **Author Contributions:** AWA and PC obtained, prepared, and cryocooled oocytes. AWA, RET, and MJM collected X-ray diffraction and image data. DWM, AWA, and RET analyzed and modelled the diffraction data. RET and SHC wrote the manuscript. **Competing Interest Statement:** RET is founder, Chairman and CTO of MiTeGen, LLC, in which he has a significant financial interest. MiTeGen manufactures and distributes tools and instruments for cryo-crystallography and cryo-electron microscopy. These include the sample supports, cryocrystallography tools, and the automated cryocooling instrument used in this work. The sample supports and automated cryocooling instrument are based on intellectual property licensed from Cornell University that was developed in RET’s Cornell laboratory. **Classification:** Major 1: Biological sciences. Minor 1: Applied Biological Sciences Major 2: Physical Sciences. Minor 2: Biophysics and computational biology.

## Abstract

Damage from ice and potential toxicity of ice-inhibiting cryoprotective agents (CPAs) are key issues in assisted reproduction using cryopreserved oocytes and embryos. We use synchrotron-based time-resolved x-ray diffraction and tools from protein cryocrystallography to characterize ice formation within bovine oocytes after cooling at rates between ∼1000 °C/min and ∼600,000°C /min and during warming at rates between 20,000 and 150,000 °C /min. Maximum crystalline ice diffraction intensity, maximum ice volume, and maximum ice grain size are always observed during warming. All decrease with increasing CPA concentration, consistent with the decreasing free water fraction. With the cooling rates, warming rates and CPA concentrations of current practice, oocytes may show no ice after cooling but always develop substantial ice fractions on warming, and modestly reducing CPA concentrations causes substantial ice to form during cooling. With much larger cooling and warming rates achieved using cryocrystallography tools, oocytes soaked as in current practice remain essentially ice free during both cooling and warming, and when soaked in half-strength CPA solution oocytes remain ice free after cooling and develop small grain ice during warming. These results clarify the roles of cooling, warming, and CPA concentration in generating ice in oocytes, establish the character of ice formed, and suggest that substantial further improvements in warming rates are feasible. Ice formation can be eliminated as a factor affecting post-thaw oocyte viability and development, allowing other deleterious effects of the cryopreservation cycle to be studied, and osmotic stress and CPA toxicity reduced.

**Significance Statement:** Cryopreservation of oocytes and embryos is critical in assisted reproduction of humans and domestic animals and in preservation of endangered species. Success rates are limited by damage from crystalline ice, toxicity of cryoprotective agents (CPAs), and damage from osmotic stress. Time-resolved x-ray diffraction of bovine oocytes shows that ice forms much more readily during warming than during cooling, that maximum ice fractions always occur during warming, and that the tools and large CPA concentrations of current protocols can at best only prevent ice formation during cooling. Using tools from cryocrystallography that give dramatically larger cooling and warming rates, ice formation can be completely eliminated and required CPA concentrations substantially reduced, expanding the scope for species-specific optimization of post-thaw reproductive outcomes.

## 1. **Introduction**

Effective cryopreservation^1–3^ of oocytes^4–6^ and embryos^7^ is critical for routine application and mass adoption of assisted reproduction technology for humans^8^, for propagation of domestic^9–11^ and endangered^12–14^ animals, and for long-term preservation of valuable genetics. Overall reproductive success rates using frozen oocytes and embryos relative to unfrozen controls vary widely between species and can vary widely between laboratories and practitioners.

The mechanisms by which cryopreservation protocols damage cells and degrade reproductive outcomes are incompletely understood, but many are related to ice formation and its inhibition. Ice may nucleate when the sample temperature drops below the equilibrium melting temperature *T_m_*, and the nucleation rate becomes very large near the homogeneous nucleation temperature (*T_h_* ∼ 236 K in pure water)^15,16^. Ice crystal growth rates in aqueous CPA solutions peak at temperatures between *T* and *T*, and become negligible at cryogenic temperatures.^17^ To inhibit ice formation, oocytes/embryos are soaked in equilibration and vitrification solutions containing penetrating (e.g., ethylene glycol (EG), dimethyl sulfoxide (DMSO)) and non-penetrating (e.g., sucrose) cryoprotective agents (CPAs) to reduce the “free” (non-hydration) water^18^ concentration; cooled to cryogenic temperature; stored; and then thawed in solutions supporting re-expansion and growth. Penetrating CPAs may be toxic, especially at high concentrations^19–22^. Ice crystals formed inside cells may puncture membranes and disrupt the spindle apparatus, cytoskeleton, and other cellular structures. Ice crystals are nearly pure water, so their growth concentrates proteins, other cytoplasmic solutes, and CPAs in remaining uncrystallized solution, which may lead to protein aggregation and denaturation and increase CPA toxicity. Physicochemical properties including solubility, pH, pKas of amino acids, and the hydrophobic effect are all temperature dependent^23^. Even without ice formation, these may cause conformational changes, partial unfolding, and aggregation of proteins^24^ and changes in structure of other cellular components, some of which may be irreversible. Ice formation and differential contraction of sample regions during cooling can cause sample fracturing – especially in larger samples – that disrupts membranes and other cellular structures.^25–28^ Oocyte and embryo cryopreservation has used two approaches,^2^ the first with cooling rates of ∼0.1-10 °C /min and the second with cooling rates greater than ∼5000 °C /min. In the first approach, ice formation inside cells is (ideally) eliminated by a combination of slow cell dehydration via nucleation and growth of extracellular ice and by modest concentrations of penetrating cryoprotectants. In the second approach, cell dehydration during cooling is negligible, and soaks in solutions containing large non-penetrating CPA concentrations (e.g., 0.5 M sucrose) to dehydrate cells and large penetrating CPA concentrations (e.g., ∼15% DMSO and ∼15% ethylene glycol) are required to inhibit ice formation in the remaining internal solvent. To minimize damage by osmotic shock, cells are soaked in a series of solutions with increasing CPA concentration^4^. Bovine oocytes have poor development outcomes after cryopreservation using the first approach but have acceptable development outcomes using the second, fast cooling approach. Oocytes are significantly more vulnerable to cryodamage than embryos, in part because oocytes are much larger, have very different internal structure, and are more primed for apoptosis, and because embryos are more primed for cell proliferation and can survive even if some of their cells are damaged and die.

Experiments by Seki, Mazur and coworkers suggest that warming is more important than cooling in determining oocyte and embryo survival and reproductive outcomes.^29–34^ During warming, water molecules initially locked in a vitreous state become increasingly mobile. Pre-existing crystallites can grow, new crystals can nucleate and grow, and larger crystals can grow at the expense of smaller ones (recrystallization).

Ice within cells can be visually detected as darkness or milkiness. This optical assay is neither sensitive nor quantitative, especially when applied to cells (e.g., bovine oocytes) with dark, lipid-rich cytoplasm. Cryocooled cells can be examined by transmission electron microscopy and ice crystals directly imaged^35^. Rates of ice formation and amounts of ice formed during cooling and warming can be assessed using calorimetry^36–38^ if cooling and warming rates are sufficiently small. Studying ice formation in oocytes during rapid warming is particularly challenging as observations must be made during the short warming transient, requiring time resolutions <<100 ms.

Synchrotron-based x-ray diffraction provides a powerful and quantitative approach to studying ice in cryopreserved samples^39,40^. X-rays easily penetrate through bovine oocytes, which are similar in size (∼100 μm) and x-ray absorption coefficient to protein crystals used in biomolecular crystallography^41,42^. X-ray diffraction can detect ice volume fractions well below 1% and the total ice diffraction flux is proportional to the amount of ice present. X-ray diffraction has been widely used to study ice formation in other contexts, including in aqueous cryoprotectant solutions and in protein crystals^39,43^. X-ray diffraction has been used to observe ice in cryocooled embryos, cumulus-oocyte complexes, and morulae^40^, but has not been systematically applied.

Here we report a comprehensive study of ice formation in bovine oocytes using static and time-resolved synchrotron-based x-ray diffraction. Diffraction data collected from cold samples is used to characterize the amount and structure of ice as a function of cooling rate, the concentrations of penetrating and non-penetrating cryoprotectants, and the type of sample support. Time-resolved diffraction data collected as these samples are warmed is used to characterize how the structure and amount of ice changes through to final melting. Much larger cooling and warming rates achieved using sample supports and a cooling instrument developed for cryocrystallography greatly reduce ice formation and the cryoprotectant concentrations required to inhibit it and allow cryopreservation without ice formation. Similarities with the bovine model^44^ suggest that our methods for optimizing cryopreservation should be applicable to human oocytes and embryos.

## Results

Oocytes were placed either on crystallography supports or on Cryotop supports popular in assisted reproduction, and cooled at ∼30,000 °C/min or ∼600,000 °C/min (when using crystallography supports) in liquid nitrogen (LN_2_) using an automated cryocrystallography instrument (**Section S3**). During x-ray data collection, these samples were kept cold using a *T*=100 K N_2_ gas stream. To warm the samples *in situ*, the *T*=100 K gas stream was blocked using air blade shutter and a room temperature N_2_ gas stream directed at the sample (**Figure S1**).

### Diffraction patterns and ice observed after cooling

#### Types of diffraction patterns observed

Observed 2D diffraction patterns from bovine oocytes at *T*=100 K vary with CPA concentration and cooling rate. As shown in **Figures 1, 2** and **S5**, the patterns are of five basic types: (1) two broad, diffuse and azimuthally uniform rings, one near 3.7 Å and a weaker ring near 2.2 Å, consistent with low-density amorphous ice *I_LDA_*; (2) three broad, diffuse and azimuthally uniform rings at expected cubic ice *I_c_*resolutions, consistent with a small-grain powder of stacking disordered ice *I_sd_*^45,46^ (**Section S6**, **Figure S6**) with a small hexagonal plane fraction *Φ*_h_; (3) four or more narrower, azimuthally continuous and “smooth” rings at resolutions expected for *I_c_* and *I_h_*, consistent with a small grain powder of *I_sd_* with a larger hexagonal plane fraction *Φ*_h_; (4) sharp, continuous rings with significant azimuthal variation, consistent with largely hexagonal ice *I_h_* and the presence of some larger grains; and (5) multiple discrete peaks on top of weaker, azimuthally continuous but nonuniform rings, indicating the presence of a modest number of even larger *I_h_*, crystals (**Figure S5**). For all CPA concentrations and cooling rates used here, diffraction as in (5) was observed only when an error allowed partial thawing and then refreezing of the sample. Measurements using ooyctes that were moved through oil before mounting and cooling (**Section S2**) confirmed that observed diffraction arose from ice inside the oocyte and not on its surface.

**Figure 1.**
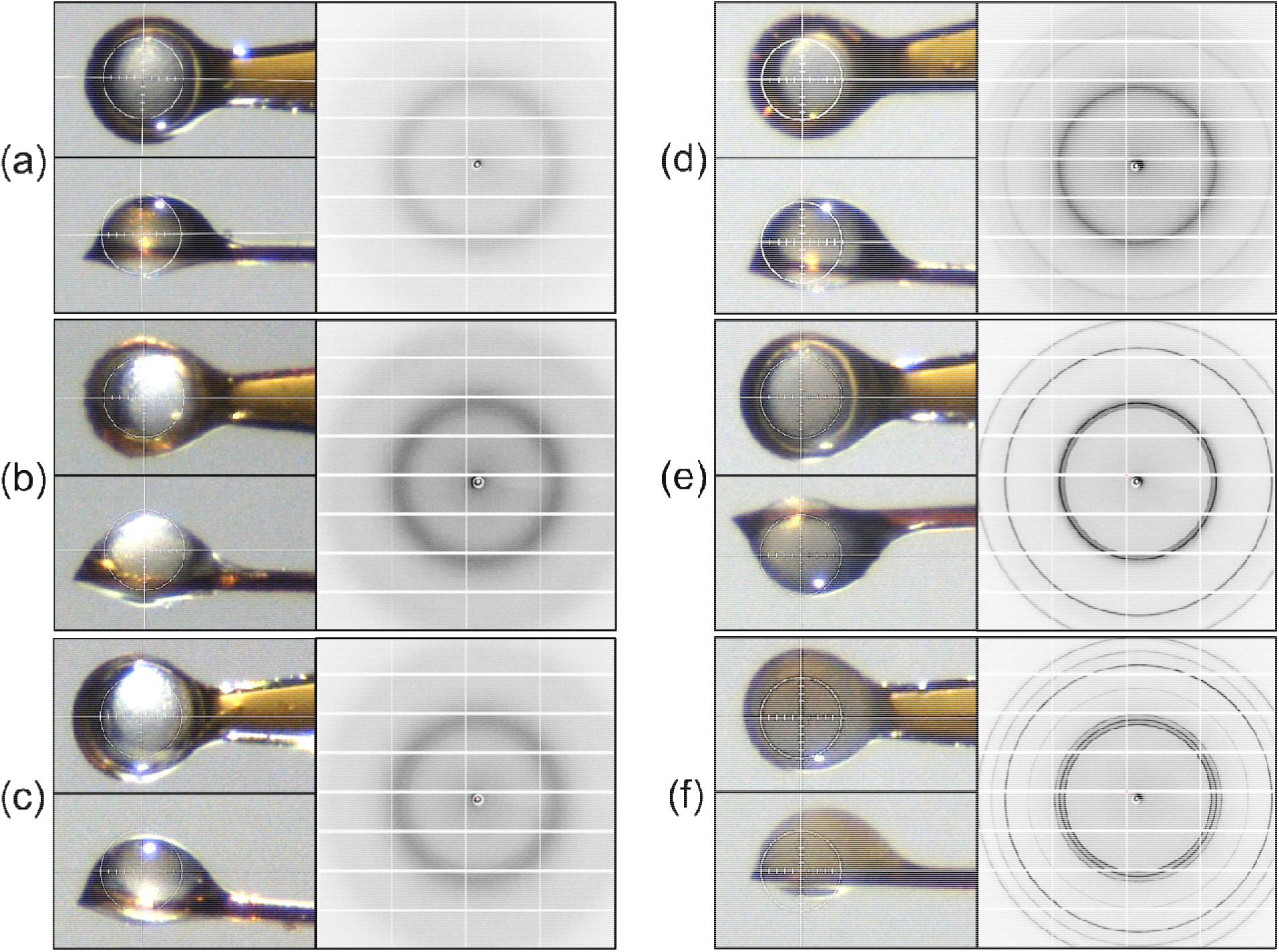
Correlation between optical images and diffraction patterns of cryocooled bovine oocytes at *T*=100 K. (a) Oocyte soaked in 15% DMSO, 15% EG., 0.5 M sucrose and “fast” cooled in LN_2_, as described in the text, showing two diffuse rings characteristic of low-density amorphous ice *I*_LDA_. (b) Oocyte soaked in 7.5% DMSO, 7.5% EG, 0.25 M sucrose and “fast” cooled. Diffuse scattering characteristic of *I*_LDA_. (c) Oocyte soaked in 13.5% DMSO, 13.5 % EG, 0.45 M sucrose and “slow” cooled in cold N_2_ gas before plunging in LN_2_. Diffuse but somewhat sharper scattering, largely consistent with *I*_LDA_. (d) Oocyte soaked in 12% DMSO, 12% EG, 0.4 M sucrose and “slow” cooled, showing well defined but somewhat broad diffraction rings at resolutions expected for cubic ice *I*_c_. (e) Oocyte soaked in 6% DMSO, 6% EG and 0.2 M sucrose and “fast” cooled. Sharp diffraction rings at cubic ice resolutions and beginning of hexagonal ice ring. (f) Oocyte soaked in 6% DMSO, 6% EG and 0.2 M sucrose and “slow” cooled. Strong and azimuthally inhomogeneous rings at all expected resolutions of hexagonal ice *I*_h_. White circle in each oocyte image has a diameter of 100 μm.

Figure 2 shows diffraction intensity versus resolution, obtained from the 2D diffraction in Figure 1 by azimuthal integration and background subtraction. The black dotted lines are best fits to a model of stacking disordered ice^46^ with the hexagonal stacking fractions *Φ*_h_ indicated (**Section S6, Figure S6**). The quality of such fits is generally excellent for all oocytes examined, particularly when the cubic stacking fraction *Φ***_c_ =**1-*Φ*_h_ is substantial. Deviations between data and fit may be due in part to background modeling and due to sample inhomogeneity (e.g., the presence of ice crystals with a distribution of *Φ*_h_ values and crystal sizes.)

**Figure 2.**
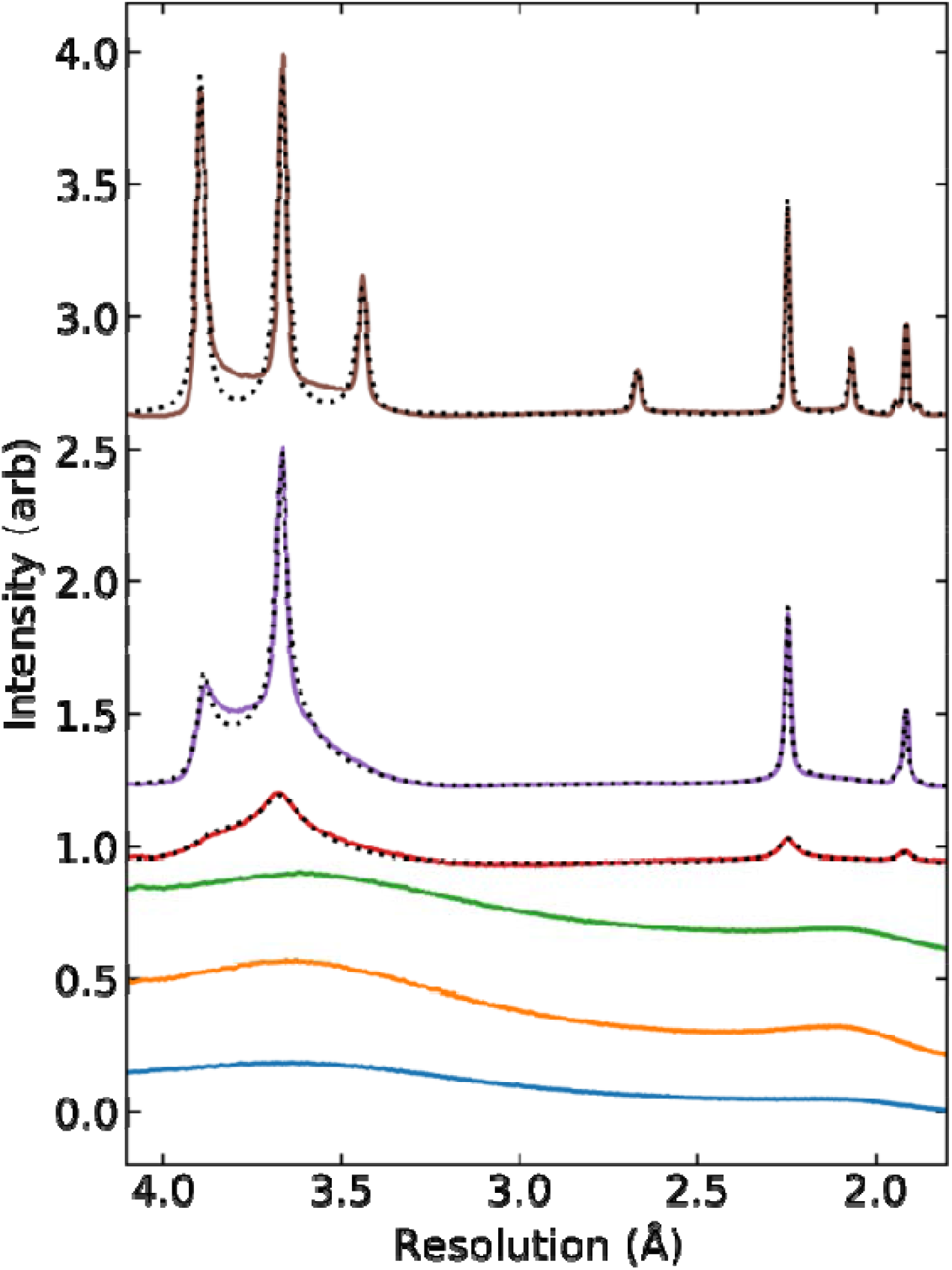
Variation of ice diffraction intensity versus resolution with amount and type of ice. Azimuthally integrated and background subtracted diffraction intensity corresponding to the samples and x-ray detector images shown in Figure 1 (a)-(f), arranged from bottom to top. The dotted black lines for the top three are fits to a model of stacking disordered ice with hexagonal plane stacking fractions *Φ* of (d) 0.35, (e) 0.40, and (f) 0.72.

#### Correlation between optical images at T=100 K and x-ray diffraction

Optical images at *T* =100 K were acquired using the beamline camera (Figure 1). When x-ray diffraction showed no evidence of crystalline ice, oocytes showed internal “grains” with an opalescent character, associated with high lipid content. When x-ray diffraction showed largely “cubic” ice (as in Figure 1(d)), optical images were largely indistinguishable from those of fully vitrified oocytes. When x-ray diffraction indicated the presence of largely hexagonal ice (Figure 1(f)), the oocytes had a milky / cloudy appearance that completely obscured the oocyte’s internal structure. X-ray diffraction produces substantial ice signals even when ice crystallites are too small or sparse to generate strong optical scattering.

#### Effects of CPA concentration and cooling rate on ice formed during cooling

Cryoprotectant concentrations required to obtain plunge-cooled oocytes with no visible crystalline ice diffraction decrease with increasing cooling rate. For oocytes soaked in vitrification solution containing 13.5% EG, 13.5% DMSO, and 0.45 M sucrose on crystallography loops and cooled at ∼30,000 °C/min, the diffraction pattern (Figure 1(b)) consisted of continuous and azimuthally uniform ice rings, consistent with powder-like ice, at the resolutions of *I*_c_. Fits to intensity versus resolution plots indicate that the ice was stacking disordered *I_sd_*with a large cubic fraction.

For oocytes on crystallography loops (**Figure S2(a)**) cooled at ∼600,000 °C/min, ice-free diffraction patterns were obtained with CPA concentrations as low as 7.5% EG, 7.5% DMSO, and 0.25M sucrose (Figure 1(b)) – the concentrations in the standard equilibration solution. At 6% EG, 6% DMSO and 0.2 M sucrose, azimuthally uniform ice rings, consistent with very small grain size ice *I*_sd_ with a large cubic fraction, were observed (Figure 1(e)). This suggests that ultrafast cooling allows the second step in the standard two-step cryoprotectant soaking protocol used for bovine oocytes to be eliminated. The factor of two reduction in minimum CPA concentration for a factor of 20 increase in cooling rate is consistent with the behavior of drops of single CPA solutions, for which the minimum CPA concentration varies logarithmically with inverse cooling rate^47^.

For oocytes mounted on thin Cryotops (**Figure S2(b)**) and cooled using plunge cooler settings that gave the largest cooling rates, no ice was observed at CPA concentrations of 12.5% EG, 12.5% DMSO, and 0.45M sucrose, and diffuse, largely cubic *I_s_*_d_ was observed at 10% EG, 10% DMSO, and 0.4 M sucrose. Oocytes on the thicker Cryotops (**Figure S2(c)**) showed strong, sharp diffraction consistent with largely hexagonal *I*_sd_ at the latter concentration. These results indicate that cooling rates achieved using Cryotops are substantially lower than those using cryocrystallography supports under otherwise identical conditions.

In all these experiments, as CPA concentrations were decreased, ice diffraction patterns followed the progression shown in Figure 1 from *I_sd_* with a large cubic fraction to *I_sd_* with a small cubic fraction to *I_h_*.

### Estimates of warming rates during warming

Oocytes were thawed *in situ* at the x-ray beamline using a room temperature N_2_ gas stream while diffraction data was recorded (**Figure S1**). Oocyte warming rates were estimated from 12 fps videos recorded using the beamline camera, from evolution of x-ray diffraction patterns, and from evolution of the ice unit cell volume (**Section S7**). For oocytes on crystallography supports that developed ice during cooling, ice diffraction began evolving ∼0.02 s after the solenoid controlling the air blade and warming gas stream flows opened and disappeared 0.08 to 0.12 s later. This gives an average warming rate between the gas stream temperature (100 K or –173 °C) and the melting temperature of ice (∼0 °C) of ∼173 °C/0.10 s ≈ 1700 °C /s or about 100,000°C/min. With the room-temperature gas stream shut off to allow warming in stagnant air, all ice diffraction disappeared after ∼0.42 s, corresponding to an average warming rate of ∼400 °C /s or ∼24,000 °C /min.

For oocytes on thin Cryotops (**Figure S2(b)**), ice diffraction disappeared after 0.22-0.31 s in the warm gas stream, corresponding to an average warming rate of ∼650 °C/s (40,000 °C/min). For oocytes on thick, curved Cryotops, ice disappeared after 0.5-0.6 s, corresponding to an average warming rate of ∼320 °C/s (19,000 °C/min). These are factors of ∼2.5 and ∼5 smaller than are achieved on crystallography loops. The physics of heat transfer in room-temperature N_2_ gas and in LN_2_ at its boiling temperature are similar. Cryotop cooling rates should thus also be ∼2.5 and ∼5 times smaller, consistent with the larger minimum CPA concentrations they require to prevent ice formation on cooling.

Oocyte warming rates can be more accurately estimated from the resolution (2θ) values of ice rings vs time (**Section S7**). These give the equivalent hexagonal unit cell volume of ice, whose temperature dependence has been accurately determined (**Figure S7**).^48^ Azimuthally integrated and background subtracted diffraction patterns (Figure 4(a)) were fit to a model of stacking disordered ice, yielding both the ice unit cell volume and the hexagonal stacking fraction *Φ*_h_ versus time (Figure 4(b).) Average warming rates oocytes on crystallography supports between ∼100 K (–173 °C) and ∼273 K (0 °C) were typically ∼ 2500 °C/s (150,000 °C/min), and maximum rates (from the slope of unit cell versus time) were ∼200,000 °C/min. These warming rates are larger than the largest reported using Cryotops of 117,000 °C /min^30^ and larger than “typical” values of 42,000 °C /min^49^. Our rates are for warming in a gas stream, whereas the Cryotop rates are for warming in a thawing solution that has far better heat transfer properties.

### Diffraction patterns and ice observed during warming

Almost all oocytes, regardless of cooling rate, CPA concentrations, sample support type, and warming gas stream velocity, developed crystalline ice during warming. The detailed behavior depended on the amount and nature of any ice formed during cooling.

For oocytes showing no evidence of ice after cooling, diffraction progressed from the appearance and sharpening of diffuse rings at cubic ice resolutions, to appearance of additional rings at resolutions of hexagonal ice, to fade-out and disappearance of all ice diffraction. In no case did an oocyte that was ice diffraction-free after cooling develop discrete ice diffraction peaks or significant azimuthal inhomogeneity in its ice rings. Average ice crystal size remained very small throughout warming at the warming rates used here so that the x-ray illuminated volume contained a very large number of crystallites. Ice ring intensity first became significant near –70°C (∼200 K), increased until ∼-20 °C, and disappeared near 0°C (with temperatures directly determined from ice diffraction as described below.) When using crystallography supports, the time interval from appearance to disappearance of ice rings was typically 40-80 ms. These quantitative observations during rapid warming can be compared with observations of mouse oocyte “blackening” in fixed temperature and slow warming experiments^50^.

Figure 3 (**a)** shows examples of azimuthally integrated diffraction intensity versus resolution measured during warming for bovine oocytes that exhibited ice diffraction after cooling. Figure 3 **(b)** shows the equivalent hexagonal ice unit cell volume, hexagonal stacking fraction *Φ*_h_, and integrated diffraction intensity in ice versus time deduced from DIFFaX fits to the data in (a). For oocytes that exhibit rings primarily at cubic ice resolutions (Figure 3 left and middle samples; **Figure S8**) before warming, no significant change is observed until the sample temperature reaches ∼180-200 K. As warming continues, additional rings at resolutions unique to hexagonal ice appear and the rings develop significant azimuthal variations in intensity, suggesting growth of larger crystals. *Φ*_h_ begins increasing at ∼180-200 K and reaches a value of 1.0, indicating that ice has evolved to pure *I*_h_, at ∼250 K. The time from the start of ice ring evolution to ring disappearance was typically 100-130 ms. The integrated intensity in ice rings increased modestly (e.g., by 20%) or remained constant even as ice transformed from “cubic” with *Φ*_h_∼0.3-4 to hexagonal with *Φ*_h_ =1, before decreasing at later times.

**Figure 3.**
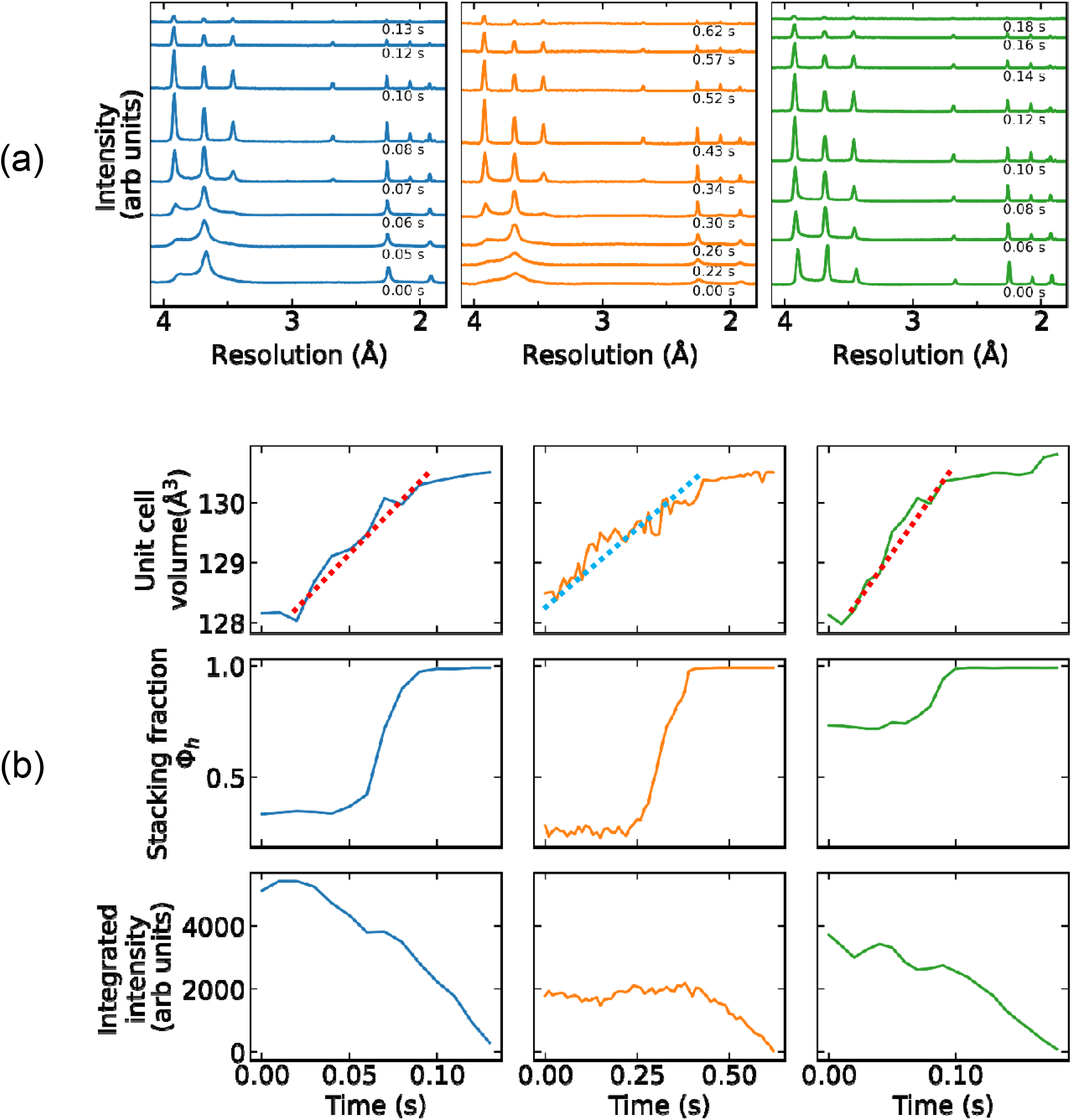
Analysis of ice diffraction during warming. (**a**) Azimuthally integrated and background subtracted diffraction intensity versus resolution and **(b)** equivalent hexagonal ice unit cell volume, hexagonal stacking fraction *Φ*_h_, and integrated diffraction intensity in ice versus time during warming of bovine oocytes. Samples from left to right: oocyte on a crystallography support, soaked in 6% DMSO, 6% EG, 0.2 M sucrose, and “fast” cooled; oocyte on a thick Cryotop, soaked in 9% DMSO, 9% EG, and 0.3 M sucrose, and “fast” cooled; and oocyte on a crystallography support, soaked in 6% DMSO, 6% EG, 0.2 M sucrose, and “slow” cooled. Parameters were extracted from DIFFaX fits to azimuthally integrated and background subtracted diffraction patterns. Time *t*=0 corresponds to the opening of the valve controlling room temperature N_2_ gas flow. Sample warming, as indicated by a change in unit cell parameter, began ∼20 ms after valve opening, due to the time required to establish warm gas flow at the sample position. Unit cell volumes of ∼128.1 Å^3^ and ∼130.5 Å^3^ correspond to temperatures of ∼100 K and ∼273 K (**Figure S7**). Dotted red lines in the upper left and right panels of **(b)** have a slope corresponding to a warming rate of ∼130,000 °C/min. The dotted turquoise line in the upper middle panel of **(b)** corresponds to a warming rate of 25,000 °C/min.

**Figure 4.**
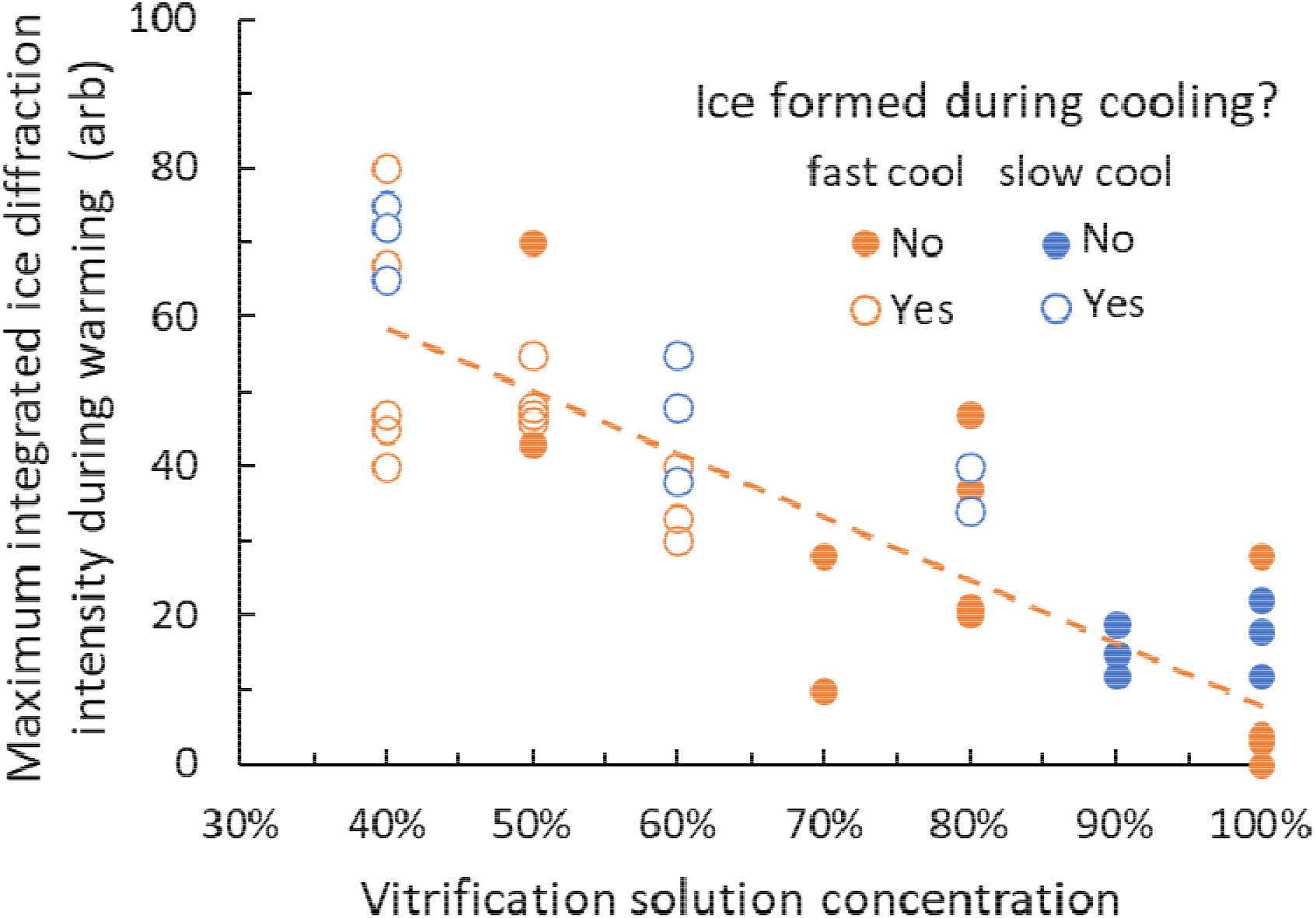
Effect of CPA concentration and cooling rate on amount of ice formed in bovine oocytes during warming. Maximum integrated intensity in ice diffraction rings observed during warming versus CPA concentration in the vitrification solution used. CPA concentration is expressed as a percentage of the standard vitrification solution used for bovine oocytes, comprised of 15% ethylene glycol, 15% DMSO, and 0.5 M sucrose. Oocytes were slow cooled (at ∼30,000 °C/min, orange circles) or fast cooled (at ∼600,000 °C/min, blue circles), and then warmed at ∼150,000 °C/min in a N_2_ gas stream. Closed and open symbols indicate oocytes that did and did not show crystalline ice diffraction after cooling and before warming. The dotted line is a fit to the fast cooled sample data. Experimental scatter may reflect variations in the initial state/quality of the oocytes, in CPA concentration within oocytes introduced in the soaks, in warm gas flow during thawing, or in the (small) amount of surface liquid (in which ice is more likely to nucleate and grow).

For oocytes with ice rings at all hexagonal positions and *Φ*_h_ >0.8 after cooling (Figure 3, right sample), the integrated intensity in ice rings and thus the amount of ice initially present was large and did not increase appreciably during warming. As the oocyte warmed above 200 K, the azimuthal “lumpiness” of the rings increased and the diffracted intensity in the brightest spots grew modestly (generally by less than ∼50%), indicating growth of modestly larger ice crystals. Larger crystallites correlated with a longer time required for all ice to finally melt: the time from the start of ice ring evolution to disappearance of the brightest spots was typically 140-220 ms.

Our data using Cryotops is more limited (**Figure S9 and S10**), but for samples that did not exhibit ice diffraction after cooling, only small grain ice was again observed during warming. Because cooling and warming times with Cryotops are longer, for given CPA concentrations more azimuthal lumpiness in the diffraction ring intensity during warming is expected. Large background diffraction intensity from the thick polymer films used in Cryotops (**Figure S9**) made analysis of ice ring behavior during warming difficult.

#### Effect of CPA concentration and cooling rate on amount of ice observed during warming

The total diffraction in all ice rings is proportional to the total amount of ice present within the X-ray illuminated volume and thus to the ice volume fraction within the oocyte. As shown in Figure 4, the maximum total diffraction in ice rings observed during warming increases roughly linearly with decreasing CPA concentration. As CPA concentration decreases, the amount of “free”, non-hydration water available for crystallization increases^18^ and maximum ice fractions should increase accordingly. Assuming additivity of the effects of EG and DMSO, data in Ref. 18 give free water fractions (by weight) in 100% and 40% strength vitrification solutions of ∼50% and ∼80%, respectively. Within the oocyte, proteins and other solutes are present in total concentrations of the same order as those in protein crystals, where 50% or more of water may be bound and unavailable for crystallization^39^. Consequently, the amount of free water within an oocyte soaked in 100% vitrification solution may be close to zero, consistent with the absence of ice formation during both fast and slow cooling. Maximum total diffraction intensities were comparable to that observed for a “snowball” oocyte in **Figure S5** (accidently thawed and then slowly cooled in the T=100 K gas stream at the x-ray beamline). This indicates that at nearly all CPA concentrations, much of the crystallizable water did in fact crystallize before final melting.

Maximum ice diffraction intensities and ice fractions observed during warming are at most only weakly dependent on cooling rate (Figure 4). Ice observed during warming is due to ice formed both on cooling and during warming, and increasing cooling rates reduces the amount of ice formed on cooling. A weak dependence on cooling rate suggests that ice forms so rapidly during warming that the maximum ice fraction is largely determined by the available crystallizable water^39^, not by how much ice nucleation and growth occurs during cooling.

### Ice free cryopreservation of bovine oocytes

Ice nucleation and growth rates decrease as CPA concentration increases. With adequate CPA concentrations and sufficiently fast cooling and warming, completing the cryopreservation cycle of cooling and warming without formation of detectable ice should be possible.

Figure 5(a) shows diffraction during warming at ∼150,000 °C/min from an oocyte that was soaked in half-strength vitrification solution (7.5% DMSO, 7.5% EG, 0.25 M sucrose) and cooled at ∼600,000 °C/min on a crystallography support. No ice diffraction is evident after cooling, and ice rings that form during warming remain continuous and homogeneous. Figure 5(b) shows diffraction during warming at ∼150,000 °C/min from an oocyte that was soaked in full-strength vitrification solution (15% DMSO, 15% EG, 0.5 M sucrose) and cooled at ∼600,000 °C/min on a crystallography support. In this case, ice diffraction that appeared during warming was extremely weak and visible for only ∼15 ms. This tiny amount of ice may have formed in residual surface liquid rather than in the solute-rich oocyte interior; extracellular ice formation typically does not affect subsequent cell survival^35,51^. Conventional warming in a thawing solution should give warming rates an order of magnitude larger and allow the CPA concentration required for an ice-free freeze-thaw cycle to be modestly reduced.

**Figure 5.**
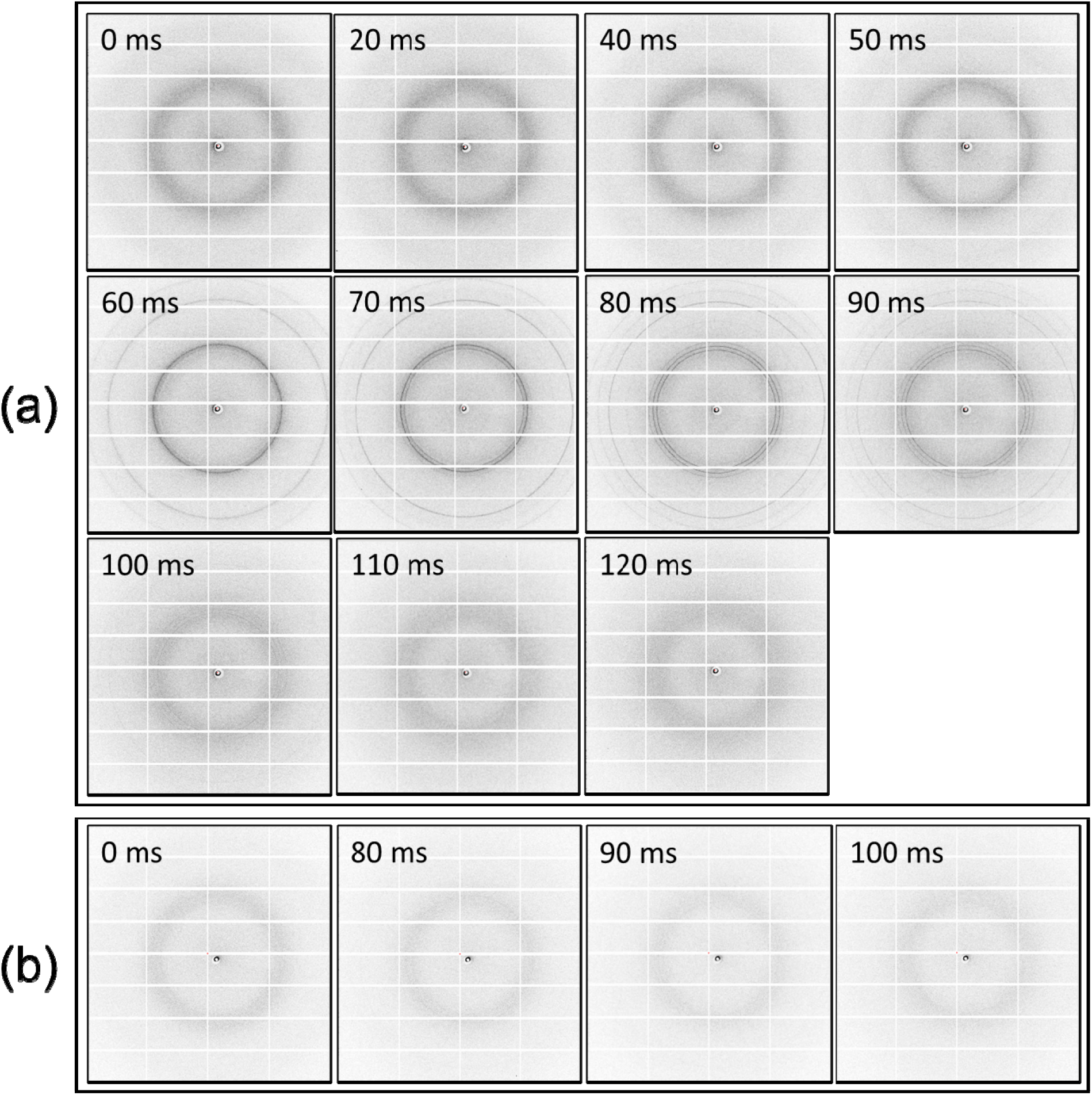
Evolution of oocyte diffraction during warming. Time series of diffraction images acquired during warming of **(a)** the oocyte of **Figure 1(b)**, soaked in 7.5% DMSO, 7.5% EG, and 0.25 M sucrose, moved through oil to remove surface solvent, and “fast” cooled; and **(b)** the oocyte of **Figure 1(a)**, soaked in 15% DMSO, 15% EG, and 0.5 M sucrose and “fast” cooled. In **(a)**, ice diffraction becomes visible at 50 ms, evolves from diffuse rings at cubic ice resolutions to sharp rings at cubic ice locations at 60 ms to rings at all hexagonal ice locations at 80 ms, and has disappeared at 110 ms. In **(b)**, ice diffraction is observed for only ∼15 ms and is extremely weak (∼4% of of (a) ring is observed and only at *t*=80 ms. **Figure S11** shows corresponding intensity vs resolution plots with DIFFaX fits.

### Estimates of ice crystallite sizes

For the cooling rates, warming rates, and CPA concentrations examined here, average ice grain sizes within bovine oocytes cooled on crystallography supports are always small compared with the ∼500 nm – 1 µm thought (based on limited evidence from a few cell types) to be large enough to cause problematic cell damage^35,52^, except in cases where oocytes were improperly cooled or accidentally thawed and refrozen at the beamline.

Methods for estimating crystal grain sizes from diffraction data are reviewed by Neher et al.^53^ When grain sizes are smaller than ∼200 nm, they can be estimated using a Scherrer/Williamson-Hall analysis of diffraction ring broadening in excess of that set by instrumental resolution (**Section S8.1**). Our large incident x-ray beam divergence of 0.2° set a maximum resolvable grain size of perhaps 30 nm. Broadening beyond instrumental resolution was only observed in diffraction patterns corresponding to largely cubic *I*_sd_ with *Φ*_h_ <0.4, either in as-cooled samples or transiently (for 10-20 ms) during warming of initially ice-free samples. In these diffraction patterns broadening due to stacking disorder is substantial and attempts to apply Williamson-Hall analysis did not yield meaningful results. No excess broadening was observed for samples with diffraction patterns corresponding to largely hexagonal ice with *Φ*_h_ >0.8.

A second bound on ice grain sizes can be obtained by estimating the minimum number of randomly oriented crystallites within the x-ray illuminated volume needed to generate continuous ice diffraction rings^39,53^ (**Section S8.2**). For our x-ray beam parameters, this would require at least ∼5000 crystals within the 10 μm diameter × 100 μm long illuminated volume. If that entire volume is crystalline ice, the average crystallite size is then ∼1.4 μm; the actual ice volume fraction was likely <0.5.

A third estimate of average ice crystallite size can be obtained by analyzing variations in the diffracted intensity with azimuthal angle χ^54^ (**Section S8.3**). In a perfect powder sample, where the x-ray illuminated volume contains a huge number of tiny, randomly oriented crystallites, the diffracted intensity will be uniform around each ring. As the number of crystallites within the illuminated volume decreases, intensity fluctuations around the ring will grow, and when only a few crystallites are present, the ring will be replaced by a handful of discrete spots.

Samples exhibiting the largest and most azimuthally nonuniform ice diffraction intensities – observed after all ice had converted to *I*_h_ and before final melting when grain sizes are largest – were selected and analysed following the approach of Yager and Majewski^54^ (**Section S8.3**). The resulting ice grain sizes were smaller than the ∼500 nm – 1 µm thought to be problematic, but the reliability of this analysis requires further investigation.

## Discussion

### Ice formation during cooling and warming

The progression of ice diffraction patterns and ice crystal form in cryocooled bovine oocytes as CPA concentration is increased – from pure hexagonal ice *I_h_* to stacking disordered ice *I_sd_*with increasing cubic character to low-density amorphous ice *I*_LDA_ – follows the general sequence observed vs CPA concentration in protein crystals^39^. The reverse progression is observed when initially ice-free oocytes are warmed. Ice diffraction patterns from oocytes exhibit asymmetric and peak/resolution dependent broadening (Figures 2 **and S6**) that is poorly fit if a mix of cubic and hexagonal crystals is assumed, but that is accurately reproduced by a model of stacking disordered ice^46^. For pure water, stacking disorder arises when ice growth occurs under conditions of deep supercooling or from amorphous ice^55^, as is expected to arise during cooling and warming rates achieved here.

CPA concentrations, cooling rates, and Cryotop sample supports used in current assisted reproduction practice appear adequate to give cold bovine oocytes that are ice diffraction and therefore ice free. During warming at rates comparable to the largest reported using Cryotops^30^, these same samples develop a substantial volume fraction of crystalline ice. The grain size of this ice – reflected by the azimuthal nonuniformity of ice diffraction rings – remains small through to melting. The impact of this transiently formed ice on post-thaw viability and development requires study, but the small average grain size makes large mechanical impacts less probable. This is consistent with observations that, compared with “fresh” oocytes not subject to cryopreservation protocols, the fraction of cryopreserved bovine oocytes that survive is modestly reduced, although developmental outcomes are more severely affected.

Using the same sample supports and cooling protocols, reducing CPA concentrations by 30% leads to ice formation on cooling and to more nonuniform ice diffraction and larger grain size ice during warming. Since EG and DMSO are efficient in inhibiting ice formation, finding alternative CPA combinations that are as or more effective in preventing ice formation at lower concentrations may be hard. Time-consuming soaking protocols to minimize osmotic shock may remain necessary when using current sample supports and cooling and warming protocols.

The relatively smooth ice rings and lack of substantial increases in azimuthal lumpiness observed during warming for nearly all samples studied here suggest that ice crystal growth via recrystallization was modest. This is consistent with the large warming rates and short times (of order 30 ms when using crystallography supports, and 75-150 ms using Cryotops) for ice growth in our experiments, and the modest ice growth velocities expected in the presence of substantial concentrations of CPAs and of native solutes (e.g., proteins) in the oocyte cytoplasm.

### Tools from cryocrystallography allow large reduction of CPA concentrations

Ice formation has long been an issue in x-ray cryocrystallography of biomolecules. The typical size range of biomolecular crystals – from a few micrometers to a few hundred micrometers – includes the size range of mammalian oocytes. The typical range of solvent contents – from ∼30% to ∼80% v/v – includes that of oocytes (before CPA soaks and dehydration). When bovine oocytes are handled and cooled using crystallography sample supports and cooling technology allow ice-free diffraction to obtained from cold samples even when the concentrations of EG, DMSO, and sucrose are all reduced by 50%. Lower CPA concentrations should allow the times for CPA soaking before cooling and oocyte re-expansion after thawing to be reduced, while reducing osmotic stress and CPA toxicity. The cooling rates achieved here of ∼600,000 °C/min via plunging in LN_2_ at its boiling temperature (77 K) are more than an order of magnitude larger than those reported for human oocytes and larger than the 23,000 °C/min reported when using Cryotops^56,57^ and the ∼35,000 °C/min obtained using thin wall quartz microcapillaries^58–60^. Use of “slushed” LN_2_ (cooling to its melting temperature) increased cooling rates in quartz capillaries to ∼250,000 °C/min^58^, and a similar increase (to ∼4 × 10^6^ °C/min) should be achievable using crystallography tools.

Successful cryoprotectant-free cryopreservation of mouse fibroblast cells by ink-jet printing ∼40 pL drops onto a cryocooled substrate has been reported^61^. Ice likely formed in the drops during cooling and certainly formed during warming (**Section S10**). Post-thaw cell viability approaching 90% is then likely due to tolerance of the chosen cells to small ice crystals. Bovine and human oocytes have much larger volumes (∼500 pL) than the fibroblast cells (∼5 pL) and drops used, maximum achievable cooling and warming rates are lower, and successful CPA– and dehydration-free cryopreservation seems unlikely. Complete elimination of CPAs may in any case be undesirable because they modulate thermal expansion^62^ and so can reduce stress and fracturing.

### Ice formation during cooling *and* warming can be eliminated

Cryocrystallography sample supports deliver larger cooling and warming rates (by factors of 2.5-5) than sample supports used in current assisted reproduction protocols. Using these supports, essentially no ice ever forms in oocytes soaked in standard vitrification solution, cooled “fast” and then warmed in N_2_ gas. Previous claims of ice-free oocyte cryopreservation have not been supported by direct measurements, let alone by an ice detection measurement as sensitive and quantitative as X-ray diffraction.

Warming rates directly measured here of 150,000 °C/min (average between 100 and 273 K) and 200,000 °C /min (maximum) for bovine oocytes on crystallography supports in a room temperature N_2_ gas stream are comparable to the largest warming rates (117,000 °C /min) reported for 50 µm thermocouples attached to Cryotops and plunged into thawing solutions^30^. Warming rates measured here in stagnant air of ∼24,000 °C/min are only 4-5 times smaller than those reported in thawing solutions. This is surprising, since aqueous thawing solutions have heat transfer properties that are vastly superior to those of N_2_ gas; they are also superior to those of LN_2_ and are predicted^63^ and measured^31^ to give correspondingly larger warming rates. Consequently, with proper protocol optimization, plunging oocytes on crystallography supports into thawing solutions should give warming rates of ∼10^6^ °C/min. When combined with the cooling rates demonstrated here, ice formation within oocytes in the freeze-thaw cycle should be completely and routinely eliminated using both standard vitrification solution concentrations and at least modestly reduced concentrations.

Alternative warming methods – such as using lasers to heat samples containing or coated with a strong absorber (e.g., carbon black, gold nanorods) ^33,64,65^ – can give larger warming rates (∼10^7^ °C/min from simulations) and have particular potential for achieving fast warming of large samples^66^. However, this approach requires substantial technical development to be practical in routine assisted reproduction practice, and CPA concentrations to avoid ice formation (roughly proportional to the logarithm of the warming rate^67^) will be only modestly reduced.

Larger warming (or cooling) rates than can be achieved using thawing solutions (or LN_2_ at 77 K) may be unnecessary for many species, except when CPA concentrations must be minimized or when CPA permeability is low. As our diffraction data shows, using standard sample supports, standard vitrification solution concentrations, standard cooling rates, and larger than standard warming rates, substantial ice forms in bovine oocytes during warming. Yet the standard tools and conditions yield adequate (but not perfect) post-thaw viability and adequate developmental outcomes in current practice. Ice need not be completely eliminated: maximum ice fractions and/or maximum ice grain sizes during the freeze-thaw cycle must be “small enough” to give desired success rates.

### Warming and cooling rates are both important

As previously noted^32–34^, much of the cryopreservation literature focuses on the importance of cooling rates in determining outcomes. However, maximum ice fractions and grain sizes always occur during thawing. An alternative suggestion, based on experiments demonstrating the importance of warming rates to mouse oocyte viability, is that cooling rates are unimportant as long as warming is fast enough^32–34^. This may be practically true in many cases but is not a sound basis for full optimization.

With sufficiently fast warming, additional ice growth beyond that formed during cooling can be minimized. In that case, whatever ice forms during cooling will determine outcomes. As long as cooling rates are larger than the critical cooling rate (CCR), the warming rate will largely determine the maximum ice fraction and grain size.

However, the critical warming rate (CWR) required for no ice formation in a nominally ice-free sample decreases as cooling rate increases beyond the critical cooling rate^67^. At the CCR, the ice fraction in the sample will be below a detection limit – perhaps 0.1% with X-ray diffraction, more when using other methods. At larger cooling rates, the ice fraction will be reduced in inverse proportion but will still be finite, providing seeds for growth during warming. As the number density of these seeds drops, the “fast enough” warming rate required to keep maximum ice fractions below a target value will drop.

### X-ray diffraction as a probe of ice and temperature

X-ray diffraction provides a sensitive probe of ice in oocytes, giving information about ice type and average grain size. With a calibrant ice sample, the integrated intensity of ice diffraction could be used to quantitatively estimate the crystalline ice fraction.

X-ray diffraction also provides a non-invasive and accurate way to measure the temperature and warming rates of ice-containing samples held in gas streams, with a detector-dependent time resolution of ∼0.1-10 ms. Oocyte cooling and warming rates are typically estimated using thermocouples having very different thermal response times or using simulations that make assumptions violated in experiment. Performing x-ray diffraction on oocytes in liquid nitrogen or in thawing solution while maximizing cooling and warming rates presents significant challenges. Instead, measurements of ice diffraction from oocytes in gas streams can be compared with thermocouple measurements in the same gas streams and in the cooling/warming liquids, allowing scaling of the sample’s gas stream cooling/warming rates to those in the liquids.

### Ice grain sizes and “black swans”

For the cooling rates, warming rates, and CPA concentrations examined here, our data and analysis suggest that average ice grain sizes are always much smaller than the 500 nm – 1 µm minimum size thought necessary for serious cell damage. However, we have no reliable information about the distribution of ice grain sizes within the oocytes (**Section S9**).

Depending upon the grain size distribution, there may be an appreciable probability of damagingly large ice crystals occurring somewhere within the oocyte volume. There may also be “black swans” – ice crystals that, based on the measured distribution, are improbably large^68^ – that could play an outsize role in determining whether an oocyte survives and thrives. These may be associated with inhomogeneities within the oocyte, e.g., cellular compartments with different internal compositions/concentrations, solvent rich / CPA poor pockets including those that form as vitrified solvent crystallizes, and interior regions disrupted by handling, ice crystal growth, or microfracturing.

### Summary and future prospects

Ice formation during cooling and warming has long been considered a key issue in cryopreservation of oocytes and embryos for assisted reproduction. Synchrotron-based x-ray diffraction provides a robust, straightforward, and high-throughput method to characterize ice inside oocytes and how its formation and properties depend on factors including cooling rate, warming rate, and CPA concentration. Substantial ice is observed during warming even in samples that show no ice diffraction after cooling and, in all cases, the largest ice diffraction intensities and ice grain sizes are observed during warming. Using tools from cryocrystallography, bovine oocytes can be cooled and warmed sufficiently fast so that no ice ever forms, and significant further increases in both cooling and warming rates and reductions in CPA concentrations are feasible.

To the extent that ice formation, CPA toxicity, and osmotic stress are important for oocytes from a given species, the present results and methods promise to yield substantial improvements in post-thaw survival and developmental outcomes. They will allow ice formation to be ruled out as a cause of differences in outcomes between cryopreserved and fresh oocytes, allowing other damage mechanisms to be identified. The principles and methods demonstrated here should be broadly applicable in optimizing cryopreservation of small biological samples.

## Materials and Methods

### Oocyte preparation and cryopreservation

Denuded MII stage bovine oocytes were prepared as described in **Section S1**. Standard equilibration solution (ES) used in cryoprotection of bovine oocytes contain 7.5% DMSO and 7.5% EG and standard vitrification solution (VS) contains 15% v/v DMSO, 15% v/v EG, and 0.5 M sucrose. Oocytes were soaked in ES and VS concentrations ranging from 40% to 100% of these standard values, as described in **Section S2**.

Immediately after the vitrification solution soak, oocytes were placed with as little surrounding vitrification solution as possible onto one of three polymer sample supports **(Figure S2)**: a 10 μm thick cryocrystallography sample support with a 100 μm aperture (MicroLoop LD 100, MiTeGen, Ithaca, NY), a flat, ∼90 μm thick Cryotop (Kitazako, Japan), commonly used in human and animal oocyte and embryo cryopreservation, or a curved, 200 μm thick Cryotop. To facilitate handling at the synchrotron, the sample supporting end of each Cryotop was removed and inserted into a cryocrystallography goniometer base. To verify that observed ice diffraction originated from within the oocyte, some were translated through a low-viscosity oil to remove aqueous surface solution.

Oocytes on supports were immediately loaded into an automated cryocooling instrument developed for cryocrystallography (**Section S3**, **Figure S3**), plunged into liquid nitrogen (LN_2_) at 77 K, and automatically loaded into 16-sample cryocrystallography “pucks” (**Figure S4**). The automated cryocooler can be operated in several modes that give different cooling rates. Cooling rates in “fast” mode for 25 μm junction bead thermocouples are >3,000,000 °C/min. Cooling rates for 100 μm diameter bovine oocytes on cryocrystallography supports are estimated to be ∼600,000°C /min in “fast” mode and ∼30,000 °C /min in “slow” mode (**Section S3**). The “slow” value is comparable to the largest “open system” cooling rates (i.e., where the ooycte/embryo directly contacts LN_2_) achieved in routine IVF practice^49^ and roughly half the cooling rate of ∼69,000°C/min^31^ reported for a 50 μm thermocouple attached to a Cryotop.

### X-ray data collection and oocyte warming

X-ray diffraction data was collected from a total of 179 bovine oocytes at MacCHESS beamline ID7B2 at the Cornell High-Energy Synchrotron Source (CHESS) using the experimental setup shown in **Figure S1**, as described in **Section S4**. A 12.8 keV x-ray beam focused to a 10 × 10 μm (FWHM) spot was directed through the oocyte center, and x-ray diffraction was recorded using a Pilatus 6M detector framing at 100 Hz, giving 10 ms time resolution. Oocytes were cooled by a *T*=100 K (–196 °C) dry N_2_ gas stream, and warmed by simultaneously interrupting the cold stream using a room temperature N_2_ gas “air blade” and initiating a flow of room temperature N_2_ gas at the sample.

### Processing and modelling of ice diffraction

Diffraction patterns captured by an area detector record the scattered x-ray intensity at different angles 2*θ* relative to the incident x-ray beam direction, which by Bragg’s law correspond to different resolutions *d* = *λ* / 2 sin(*θ*).

Processing and modeling of detector frames and ice diffraction largely followed the protocol used by Moreau et al.^39^ in a study of ice diffraction from protein crystals. Detector frames were processed to fit and remove background and azimuthally integrated (**Section S5**). Diffraction intensity versus resolution plots for each frame from each oocyte were analyzed (**Section S6)** by embedding the program DIFFaX^69^, which calculates diffraction from samples containing stacking faults, in an optimization routine to determine best-fit parameters for stacking disordered ice formed of planes of hexagonal (*I_h_*) and cubic (*I_c_*) ice. This fitting procedure yielded four parameters for each frame / time point: the hexagonal stacking fraction *Φ*_h_, the lattice constants *a* and *c* of the equivalent hexagonal unit cell, the overall scale of the intensity, and an instrumental broadening parameter. These were then plotted versus frame number / time.

## Acknowledgments

This work was supported by Agriculture and Food Research Initiative Competitive Grant no. 2022-67015-37225) from the USDA National Institute of Food and Agriculture, and by the Cornell University Center for Advanced Technology (CAT) FY2019-2020. MJM and DWM acknowledge support by the National Institutes of Health (NIH) under award 5R01GM127528-04. Development of the cryocooling instrument was supported by the NIH under award 5R44GM101817-03. This work is based upon research conducted at the Center for High Energy X-ray Sciences (CHEXS), which is supported by the National Science Foundation under award DMR-1829070, and at the Macromolecular Diffraction at CHESS (MacCHESS) facility, which is supported by NIH award 1-P30-GM124166-01A1 and by New York State’s Empire State Development Corporation (NYSTAR). B. Miller, M. Szebenyi, and D. Schuller, and I. Kriksunov provided assistance at CHESS. Y. L. Lee, L. H. Aguiar and M. Diel de Amorim provided technical support for embryology and media preparation. David Closs assisted with the automated cryocooling instrument. RET acknowledges Brian Wowk for helpful discussions.

