## Supporting Information for "Ice formation and its elimination in cryopreservation of bovine oocytes"

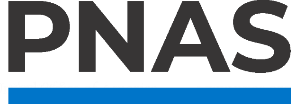

**Supporting Information for**

**Ice formation and its elimination in cryopreservation of bovine oocytes**

Abdallah W. Abdelhady, David W. Mittan-Moreau, Patrick L. Crane, Matthew J. McLeod, Soon Hon Cheong, Robert E. Thorne

**Corresponding authors:** Robert E. Thorne, Soon Hon Cheong

**This PDF file includes:**

Supporting text

Figures S1 to S12

Tables S1 to S2

SI References

Supporting Information Text

S1. Preparation of denuded MII stage bovine oocytes

Bovine ovaries were obtained at a slaughterhouse (Cargill, Wyalusing, PA), washed in 0.9% saline solution, and then transported to our laboratory (two hours by car from the slaughterhouse) while held at temperatures between 22-27 °C. Before aspiration, ovaries were washed several times in warmed saline solution. Cumulus-oocyte complexes (COCs) were aspirated from follicles of size between 2 and 8 mm using an 18-gauge needle attached to an aspiration unit consisting of a vacuum manifold and collected in 50 mL conical tubes. Good quality COCs defined as those with a dark homogeneous cytoplasm and surrounded by 3-4 intact layers of cumulus cells were selected for maturation. COCs were matured in IVM media (BO-IVM, IVF Biosciences, UK) within 5-well plates. Each well contained up to 50 COCs in 700 μL of media. COC-containing plates were incubated at 38.5 °C for 22 hours. Matured COCs were denuded by vortexing at 3200 rpm for 30 s in 50 μL BO-Wash. MII stage oocytes were selected based on the presence of the first polar body and transferred to a holding medium comprised of M-199 (Sigma M2520, USA) with Hank’s balanced salts containing HEPES (25.04 mM) and L-glutamine (0.685 mM) (Gibco, USA) supplemented with 10% fetal bovine serum, 0.2 mM sodium pyruvate, gentamycin (25 μg/mL) and pH balanced to 7.3 ± 0.1.

S2. Soaking protocol for cryoprotection of oocytes

Standard equilibration solution (ES) used in cryoprotection of bovine oocytes consists of the same base media M-199 used in the holding media with supplementation of 7.5% v/v DMSO and 7.5% v/v EG and standard vitrification solution (VS) contains 15% v/v DMSO, 15% v/v EG, and 0.5 M sucrose. Oocytes were soaked in ES and VS concentrations ranging from 40% to 100% of these standard values, as shown in **Table S1.** Immediately prior to cryocooling, each oocyte was incubated for 60 s in basic solution (BS) comprised of 100-μl PBS supplemented with 20% FBS, held on a 65 mm diameter glass dish. Three separate 100 μl drops of equilibration solution (ES), containing DMSO and EG at half the final target concentration in BS, were placed on the dish adjacent to the BS drop. Under a stereomicroscope, the BS drop containing oocytes was gently connected to the nearest ES drop. After 60 s the BS-ES drop was connected to the second ES drop, and then after 60 s this was connected to the third ES drop. The equilibrated oocyte was transferred to and incubated in vitrification solution (VS), which contained DMSO and EG at the target concentration as well as sucrose in BS, for 20 to 25 s, and then loaded either onto a crystallography support (**Figure S2(a)**), a “thin” Cryotop™ IVF support (**Figure S2(b)**), or a thick Cryotop™ IVF support (**Figure S2(c)**). The cryocrystallography supports are 10 μm thick and support the sample in a circular loop. They minimize thermal mass, maximize access of cooling and warming media to the sample, and minimize background x-ray scatter from the support. Current and original Cryotops support the sample on a flat ~90 μm thick polymer sheet and on a curved ~200 μm thick polymer sheet, respectively.

Oocytes were transferred onto Cryotops using a pipette, and onto crystallography supports by “scooping” them from a shallow concave well, in all cases leaving as little surrounding vitrification solution as possible ^1^.

Ice may form both inside the oocyte and in residual vitrification solution on its surface. Since concentrations of all solutes including CPAs inside the oocyte differ from those in surface solution, the amount and character of ice formed is expected be different. Crystallography loops minimized the amount of surrounding solution (**Figure 1)**, so that the volume of surface liquid illuminated by the 10 µm x-ray beam was a small fraction of the illuminated volume of the oocyte. To further rule out surface liquid as a significant source of observed ice diffraction, oocytes soaked in the full range of CPA concentrations explored here were transferred to and moved within a low-viscosity oil (LV CryoOil, MiTeGen) (used in cryocrystallography to remove aqueous surface solvent from protein crystals) to remove surface solution and then placed on crystallography loops. Diffraction results obtained at each CPA concentration appeared to be more reproducible, but otherwise were qualitatively and quantitatively similar to those obtained without oil treatment. This confirmed that any ice formed in surface solution for well-prepared samples did not contribute significantly to observed diffraction.

S3. Oocyte cryocooling, storage, transport, and handling at the synchrotron source

Oocytes on supports were plunged into liquid nitrogen (LN_2_) at 77 K and then automatically loaded into 16-sample cryocrystallography “pucks” (Universal V1 Puck, Crystal Positioning Systems) using an automated plunge cooling system developed for cryocrystallography (**Figure S3**, NANUQ^TM^, MiTeGen). The pucks were manually inserted into storage canes (Shelved Puck Shipping Cane, MiTeGen) held inside cryogenic Dewars for storage and transport to the synchrotron x-ray source **(Figure S4)**.

The automated plunge cooler can be operated in several modes with different plunge speeds, with cold gas trapped or completely removed from above the LN_2_, and with plunge bore heaters on or off. Most samples were “fast” cooled using the maximum plunge speed (2 m/s) with bore heat on and cold gas removed; some were “slow” cooled using a plunge speed of 0.04 m/s, with no bore heat and no cold gas removal. Cooling rates for oocytes were estimated based on measured cooling rates of thermocouples with different bead diameters, and by scaling these values to account for the different size and thermal properties of oocytes based on previous analysis and modeling of heat transfer^2^. The heat capacity per unit volume (not mass) of metals is similar to that of water; the heat transfer rate for samples in this size range is limited by the thermal boundary layer in the cooling fluid, not the sample’s thermal conductivity; and the cooling rate varies with sample diameter *D* roughly as *D*^3/2^.

For oocytes mounted on crystallography supports and plunged at 0.04 m/s through several cm of cold N_2_ gas before entering LN_2_, the cooling rate was of order 30,000 °C/min. For oocytes mounted on crystallography loops and plunged at 2 m/s through room temperature N_2_ gas into LN_2_, the cooling rate was of order 600,000 °C/min.

Minimum CPA concentrations to prevent ice formation in aqueous solutions vary logarithmically with inverse cooling rate^3,4^. Consequently, precise knowledge of cooling rates is not essential to determining overall behavior, provided cooling rates range over at least an order of magnitude.

All oocyte equilibration and vitrification soaks, transfer to sample supports, and transfer to the plunge cooler were performed in an environment humidified to ~80% r.h. (**Figure S3**). Combined with fast sample transfers, this ensured consistent sample hydration.

S4. X-ray data collection

X-ray data was collected in February 2021 and April 2022 using MacCHESS beamline ID7B2 at the Cornell High-Energy Synchrotron Source (CHESS). The 12.8 keV x-ray beam was focused using a compound refractive lens to 10 × 10 μm (FWHM). X-ray diffraction was recorded using a Pilatus 6M detector framing at 100 Hz, giving 10 ms time resolution. Crystallography supports and Cryotops were oriented with the plane of the polymer film perpendicular to the x-ray beam, and the beam directed through the center of the oocyte. Using a small x-ray beam maximized the ratio of x-ray scatter from the oocyte to scatter from surrounding liquid and polymer, and allowed bits of frost/ice that accumulate in LN_2_ and sometimes become attached to the oocyte surface to be avoided.

Oocyte samples in Unipucks were removed from the storage Dewar and storage canes and loaded into the LN_2_-filled chamber of the beamline sample automounter. Each sample was transferred by the automounter onto a magnetic sample rotation stage, where it was cooled by a *T*=100 K (-196 °C) dry N_2_ gas stream (Oxford Cryosystems Cryostream 700). During the transfer, samples were enclosed in a cryocooled stainless steel gripper that prevented warming. To thaw the sample, a dry room temperature N_2_ gas “air blade” interrupted the flow of cold gas and a stream of dry room-temperature N_2_ gas was directed at the sample. Both room temperature N_2_ flows came from the same source and were switched on and off by the same solenoid valve under computer control. The thawing N_2_ stream diameter was 2 mm at its outlet and expanded as it traveled ~2-3 cm (at a speed of order 1 m/s) to the sample position. **Figure S1** shows the experimental setup at the beamline.

Each sample was optically imaged at orientations of 0° and 90°, and a single x-ray diffraction frame with 0.1 s exposure acquired to verify that the sample was in the beam. Then, 210 x-ray frames, each with 0.01 s exposure, were acquired under computer control. The first 10 frames were acquired with the air blade and thawing gas streams off. These recorded diffraction from any ice formed on cooling and verified that sample motion in the *T*=100 K cryostream was negligible. After the 10^th^ frame, the air blade shutter and room temperature gas stream were switched on, and the sample warmed to room temperature while the remaining 200 frames were acquired.

Sample behavior during warming was optically recorded using the beamline camera with a frame rate of ~12 Hz. For most experiments, the warm gas stream flow rate was adjusted to near a maximum value beyond which visible deflection of the sample support and/or displacement of the thawed sample on the support were observed. During warming, the sample support (MicroLoop or Cryotop) expands, so that the center of the oocyte moves a fraction of its diameter out of the x-ray beam. This causes a small drop in illuminated sample volume and in recorded diffraction intensities, making it difficult to quantitatively estimate the evolution of ice fraction within the sample versus time during thawing. Although the warming gas stream has no moisture, the vapor pressure of ice is very small at all temperatures, so oocyte dehydration during warming was negligible until the sample melted; after melting the sample dried out in roughly 10 s. Warming rates were determined from analysis of x-ray diffraction as discussed below.

Frost – floating in the LN_2_ filled chamber of the beamline sample automounter or condensed from moisture in surrounding air onto the sample and support – sometimes adhered to the oocyte surface. Diffraction from frost was always consistent with purely hexagonal ice (*Φ*_h_=1), allowing it to be distinguished from internal ice, and always consisted of discrete, sharp spots, indicating much larger ice grain sizes than were present inside the oocyte.

S5. X-ray diffraction processing

Processing and modeling of detector frames and ice diffraction largely followed the protocol used by Moreau et al.^5^ in a study of ice diffraction from protein crystals.

For a single crystal, diffraction “spots” are observed at well-defined *d* (2*θ*) values, and at well-defined azimuthal angles *ϕ* that depend on crystal orientation. For an ideal powder comprised of a large number of randomly oriented crystallites within the illuminated x-ray volume, diffraction from each crystallite occurs at a different set of azimuthal angles *ϕ*, so the diffraction pattern consists of azimuthally uniform and continuous rings at angles 2*θ* set by the single-crystal *d* values. For a small number of larger, randomly oriented crystals, multiple diffraction spots will be observed at a given *d*, spread out azimuthally around the “powder ring”, with gaps between them. For an intermediate number of crystallites, the “powder rings” will be continuous but the intensity within the ring will vary with *ϕ*.^6^

Diffraction images were loaded into python using the package *FabIO* ^7^, which allowed the data to be handled as a numpy array, and were azimuthally averaged using the *pyFAI* integration package ^8^. *pyFAI* reduced the 2-dimensional Bragg-peak-masked diffraction data from the detector (counts at pixel coordinates *x*, *y*) to 1-dimensional diffraction data (average counts in a bin of angular width
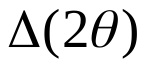
) versus angle
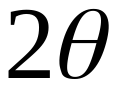
 or resolution
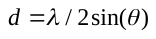
, correcting for beam polarization (90%, in the horizontal direction) and for variations in solid angle recorded by each pixel with
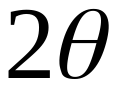
.

The non-crystalline ice x-ray background had contributions from the sample — scattering from solutes and structural components of the oocyte/embryo, from unfrozen solvent (water) including surface solvent, from oil (when used), from low-density amorphous ice, from the sample support (especially when Cryotops were used) — and from the experimental set-up (primarily air scattering). The background was determined by evaluating a 10th order polynomial fit to the difference
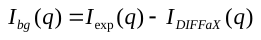
 between the *DIFFaX* model and our experimental diffraction patterns (similar to the method used in the powder diffraction analysis program *GSAS-II* ^9^), and then optimizing until the *DIFFaX* model and background converged. The resulting background fits were slowly varying curves (on the 2θ or resolution scale of the ice diffraction peaks) with a consistent shape between samples that captured the diffracted intensities from all non-crystalline sources.

S6. Modelling ice diffraction

Early measurements of ice diffraction were interpreted in terms of a mix of crystallites of two forms of ice: cubic ice *I_c_*, which formed under conditions of significant supercooling (as can occur during rapid cooling), and hexagonal ice *I_h_*. As given in **Table S2** and **Figure S6**, in the resolution range 4.0 Å to 1.8 Å, ideal cubic ice *I_c_* has diffraction peaks at resolutions of 3.66 Å, 2.25 Å and 1.92 Å; hexagonal ice *I_h_* also has peaks at these resolutions and additional strong peaks at 3.90 Å, 3.44 Å, 2.67 Å and 2.07 Å. However, observed diffraction peak shapes exhibit asymmetric broadening that is unrelated to grain size and that cannot be accounted for using a mixture of cubic and hexagonal ice crystallites. More recent measurements and analysis indicate that cubic ice *I*_c_ is almost never observed. Instead, as shown in **Figure S6**, experimental diffraction patterns are best described as arising from a stacking-disordered mix of cubic and hexagonal ice planes within each crystallite^10–12^, a conclusion supported by simulations of ice crystal growth^13,14^.

In a simple model of stacking-disordered ice *I*_sd_^10,11^ the alternation of cubic and hexagonal planes is assumed to be purely random. In this case, the ice diffraction (**Figure S6**) can be characterized by a parameter

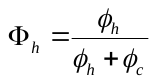
,

the average fraction of planes within each crystallite that are hexagonal.

Diffraction intensity versus resolution plots for each diffraction frame from each oocyte sample were analyzed by embedding the program DIFFaX^15^, which calculates diffraction from samples containing stacking faults, in an optimization routine to determine best-fit parameters for stacking disordered ice formed of planes of hexagonal (*I_h_*) and cubic (*I_c_*) ice. This fitting procedure yielded four parameters for each frame / time point: the hexagonal stacking fraction *Φ*_h_ the lattice constants *a* and *c* of the equivalent hexagonal unit cell, the overall scale of the intensity, and an instrumental broadening parameter. These were then plotted versus frame number / time.

In detail, observed 2D ice diffraction patterns recorded by the detector for oocytes at T=100 K varied with sample cooling rate and with the concentrations of ethylene glycol, DMSO, and sucrose. The patterns were of four basic types: (a) azimuthally uniform and diffuse scattered intensity with a single, radially very broad ring, consistent with low-density amorphous ice *I_LDA_*; (b) azimuthally uniform and diffuse scattered intensity with a two or three broad rings; consistent with very small grain size, stacking disordered ice ISD with a large cubic fraction; azimuthally lumpy, radially very narrow diffraction with intensities roughly consistent with *I_h_*; (c) azimuthally lumpy diffraction with an azimuthally uniform component, the latter component consisting of a mix of narrow and broad peaks; and (d) azimuthally uniform patterns showing a mix of narrow peaks and broad, asymmetric peaks characteristic of stacking disordered ice *I_sd_* (a disordered stacking of (001) planes of *I_h_* and (111) planes of *I_c_* ).

Azimuthally uniform ice diffraction patterns were analyzed using the program *DIFFaX* ^15^, which models diffraction from crystals containing one or more phases that may be separated by coherent planar defects including twins and stacking faults. An input file for stacking disorder was obtained from the supplementary material of Malkin *et al*.^10^. The model (**Figure S6 (c)**) consists of (0001) planes of hexagonal ice randomly stacked with (111) planes of cubic ice. The probability of a cubic plane being followed by hexagonal plane is *Φ*_ch_ and of a hexagonal plane being followed by a cubic plane is *Φ*_hc_ ^11,12^.

To optimize fits to the diffraction data, *DIFFaX* fitting and background subtraction were embedded in an optimization routine using a bounded, limited-memory BFGS algorithm implemented utilizing the *scipy.optimize.minimize* program in the *SciPy* python library ^16^. The results of this simulation were used to determine the ice crystal’s unit cell parameters, stacking probabilities, instrumental broadening, and structure factors. To verify the accuracy of the structure factors, they were also calculated using the explicit equations previously reported ^17^. As shown in Fig. 9 of ^11^, calculated ice diffraction profiles show strong variation with the stacking parameters *Φ*_ch_ and *Φ*_hc_ and thus with *Φ*_h_ so that *Φ*_h_ values obtained by fit optimization are robust. A single *B*-factor was used for the oxygens and hydrogens in the *DIFFaX* model and was fixed to 1.5, which corresponds to
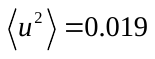
 Å^2^. The instrumental broadening was assumed to be purely Lorentzian and parameters *u*, *v*, and *w* were optimized to estimate the FWHM broadening as a function of angle.

Diffraction peaks with *I*_h_ Miller indices such that
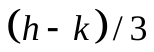
 is an integer ^10^, i.e., the (002), (110), and (112) peaks, are not broadened by stacking disorder, and the other peaks are greatly broadened. The (002) peak is overlapped by the broadened (100) and (101) peaks, the (112) peak is overlapped by the broadened (200) and (201) peaks, while the (110) peak is not overlapped.

S7. Estimates of warming rates during thawing
    S7.1. Estimates based on time for ice diffraction to disappear

For oocytes on crystallography supports, x-ray diffraction patterns showed visible evolution beginning 0.02 s after opening of a solenoid valve controlling the air blade and warming gas flows. This time delay is consistent with estimates of the time required for the air blade to interrupt the cold stream flow and for the room temperature gas jet to reach the sample. Except when large grain ice formed on cooling, all ice diffraction disappeared 0.10 to 0.14 s after opening the solenoid valve. This gives an estimate of the average sample warming rate between the gas stream temperature (100 K or -173 °C) and the melting temperature of ice (0 °C ) of ~173 °C/0.10 s ≈ 1700 °C /s or about 100,000 °C/min. With the room-temperature gas stream shut off, all ice diffraction disappeared after ~0.44 s, corresponding to an average warming rate of ~400 °C /s or ~24,000 °C /min.

For oocytes on current thin, flat Cryotops (**Figure S2(b)**), the time after opening the solenoid valve when ice diffraction disappeared was 0.24-0.33 s, corresponding to an average warming rate of ~670 °C/s (40,000 °C/min). For oocytes on previous thick, curved Cryotops (**Figure S2(c)**), ice disappeared after 0.5-0.6 s, corresponding to an average warming rate of ~320 °C/s (19,000 °C/min). These are factors of ~2.5 and ~5 smaller than are achieved on crystallography loops.

Since the physics of heat transfer in room-temperature N_2_ gas and in LN_2_ at its boiling temperature are similar, cooling rates achieved using crystallography loops should be 2.5 and 5 times larger than using Cryotops. This difference in cooling rate is consistent with our data indicating that minimum CPA concentrations to achieve no ice on cooling are substantially larger when using Cryotops than when using crystallography loops.

S7.2. Estimates based on ice unit cell volume deduced from diffraction

The equivalent hexagonal unit cell volume of ice (*I_c_*, *I_sd_*, or *I_h_*) can be determined from the resolutions (*d* values) at which ice diffraction rings are observed. Since the unit cell dimensions of hexagonal ice versus temperature have been accurately determined^18^ (**Figure S7**), *d* values and the corresponding unit cell volume provide an accurate measure of oocyte temperature. **Figure 4(b)** shows examples of equivalent hexagonal ice unit cell volumes versus time following the opening of the solenoid valve, determined using ice diffraction from bovine oocytes. Maximum warming rates (given by the maximum slopes) during thawing of oocytes on crystallography supports using an N_2_ gas stream were typically ~ 2500 °C/s (150,000 °C/min). This is larger than the average rate between -173 °C (100 K) and 0 °C (273 K) estimated above because warming rates decrease as the difference between the gas stream and sample temperatures decrease. These warming rates are comparable to the largest warming rates previously reported^19^ using Cryotops of 117,000 °C /min and larger than typical values ~42,000 °C /min^20^. However, our values are achieved using a room temperature gas stream, not by plunging into a warm aqueous solution.

S8. Estimates of ice crystallite sizes
    S8.1. Scherrer equation estimate

The size of ice crystallites formed within an oocyte can in principle be estimated using the observed ice diffraction ring breadths after azimuthal averaging. If an ice crystal has a finite size, its diffraction peaks are radially broadened by an amount inversely proportional to the crystal size. As crystal size decreases, the breadth,
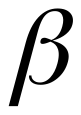
, of the diffraction peaks in
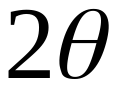
 increases according to Scherrer's equation,

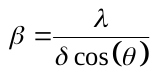
.

Here, the breadth of a peak is estimated as the integrated area beneath the peak divided by its maximum amplitude ^21^,
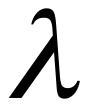
 is the wavelength,
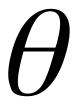
 is the Bragg angle, and
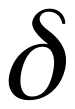
 is the apparent size of the crystallite. The actual crystallite size is proportionally related to the apparent crystallite size by
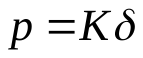
, where *K* is Scherrer’s constant ^22^. Scherrer’s constant for a cubic crystal’s (100), (110), and (111) reflections are 1.0000, 1.0607, and 1.1547 respectively ^22^. *K* is assumed to be 1 for this analysis.

Strain, dislocations, and planar faults can generate additional peak broadening ^23,24^. Limited prior knowledge of their character and distribution prevents accurate modelling of their contribution to the observed broadening ^25^. The raw measured peak breadths *β_raw_* also include contributions from instrumental broadening. This arises primarily from the dispersion and divergence of the x-ray source and from errors in defining the experimental geometry. The CHESS FLeXX beamline 7IDB2 uses a multilayer monochromator with a 0.6% energy bandpass, which gives a relatively large beam divergence.

Minimum ice crystallite sizes necessary to cause cellular damage have been suggested to be ~500 nm. Using this value for δ, our x-ray wavelength of 0.967 Å, and *θ* values for the three cubic diffraction rings of 7.59°, 12.41°, and 14.62° gives broadenings *β* of ~0.011°. The minimum measured ice diffraction peak widths were ~0.2° (corresponding to δ~30 nm) and were dominated by instrumental broadening. Consequently, peak broadening could not reliably be used to determine the sizes of crystallites larger than ~30 nm.

S8.2. Number of ice crystallites required for continuous diffraction rings

The minimum number of ice crystallites required to generate azimuthally continuous diffraction rings can be estimated from the illuminated sample volume and the incident beam divergence. This estimate can in turn be used to set a bound on the crystallite size. Assuming the ice has a sufficiently large grain size that its Bragg peaks have a breadth in the azimuthal direction determined solely by the beam divergence, α, the peak’s arc length is

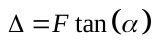
.

$F$ is the sample-to-detector distance. The circumference of the diffraction ring is

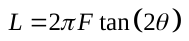
.

If placed end to end, the number of crystallites needed to generate a complete diffraction ring is
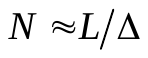
. This is multiplied by 10 to ensure there is significant overlap between the Bragg peaks to give

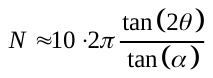
.

This corresponds to 5,000 crystallites for a diffraction peak at
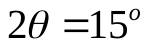
 and a beam divergence of
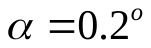
. For a 10 μm beam diameter and a 100 µm path length through an oocyte, the illuminated volume is
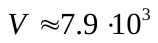
µm^3^. Dividing this total volume by 5000 gives an upper bound on the crystallite volume of 1.5 µm^3^ or, assuming a spherical crystallite, a diameter of 1.4 µm. This crystallite size gives a finite-size broadening of
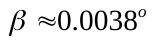
, ~50 times smaller than the observed hexagonal ice peak breadth, suggesting that the observed peak breadth is limited by the instrumental broadening, as assumed in this estimate.

S8.3. Grain size estimate from azimuthal fluctuations in ice ring intensity

A third estimate of average crystallite size can be obtained by analyzing variations in the diffracted intensity with azimuthal angle χ^6^. In a perfect powder sample, where the x-ray illuminated volume contains a huge number of tiny, randomly oriented crystallites, the diffracted intensity will be uniform around each ring; in a sample where the illuminated volume contains a small number of large crystals, each ring is replaced by a series of peaks, so the intensity around the ring is highly nonuniform. As the number of crystallites in the illuminated volume grows and their size shrinks, the fluctuations in diffracted intensity with χ relative to the mean intensity decrease.

Simulations and analysis^6^ indicate that, if the intensity *I*(χ) is regularly sampled around a diffraction ring and then these samples are replotted in order of decreasing intensity, the resulting “sorted” plot exhibits an exponential or power law decay that can be fit to obtain an estimate of the total number of crystallites within the X-ray illuminated volume. Plots of ordered pixel intensity I_sort_(χ) vs χ are fit using^6^

$I_{sort}\left( \chi\right)=a_{\psi}\exp\left( -\frac{\chi}{\psi} \right).$	**(S1)**

The decay constant *ψ* gives an estimate of the number of crystallites within the X-ray illuminated sample volume given by

$N_{g}\approx\frac{q^{3}}{4m\left( 2\pi\right)^{1/2}\sigma_{q}^{3}f_{\chi}}\psi^{2}$,	**(S2)**

where *q* is the scattering wavevector for the ice ring, *m* is the multiplicity of the ring, σ_q_ is the radial width of the ice ring, and *f*_χ_ is the fraction of the full 2π ring in χ that is sampled^6^. An upper bound on the average grain size *L* can be obtained by assuming that the entire X-ray-illuminated sample volume *V* is ice, as

$L\approx\left( \frac{V}{N_{g}} \right)^{1/3}$.	**(S3)**

In our case, *V* ≈ 10 µm × 10 µm × 100 µm. Application of this approach to samples that showed the most intense and nonuniform ice diffraction intensity (which occurred during warming before final melting) yielded grain sizes well below 500 nm. However, the estimates varied with the diffraction ring used and with frame during warming in ways that do not seem physical. The origin of these irregularities is being explored. If we cannot resolve them, discussion of this analysis will be removed from the manuscript.

S9. Limitations of our x-ray experiments

During warming, the oocyte moved laterally relative to the X-ray beam as the sample support structure holding it to the beamline’s goniometer stage thermally expanded. This changed the position and volume illuminated by the X-rays, and thus the diffracted intensities. For quantitative ice fraction estimation during warming, oocyte motion relative to the x-ray beam should be reduced by using materials for the supporting structure with low thermal expansion.

Several approaches can be used to estimate grain size distributions within “powder” samples. These typically require that the number of grains within the x-ray illuminated volume be small, that the x-ray beam divergence (the instrumental resolution) be small enough to allow the intrinsic width of ice diffraction peaks to be measured, and/or that diffraction be recorded from the sample in multiple orientatons. The large size of our x-ray illuminated volume compared with the grain size, the significant angular divergence of our x-ray beam, and our recording of diffraction from each oocyte in only a single orientation preclude meaningful modelling of grain size distributions^6,26^.

Detecting rare large ice crystals (including “black swans”) that could cause lethal damage requires collecting x-ray data from the entire oocyte. In our experiments, the 10 μm x-ray beam probed only a small fraction of the oocyte volume, and so could miss large and lethal ice crystals present elsewhere. For some cold samples at T=100 K, the oocyte was raster scanned in the beam to probe its entire volume. No change in apparent continuity of the rings was observed, and variations in intensity fluctuations within the rings (reflecting the mean ice grain size within the x-ray beam) were modest. Any large ice peaks appeared to correlate with frost on the sample surface. Large ice crystals and “black swans” may be more likely during warming, but short warming times (<100 ms) together with available raster scan speeds and maximum detector frame rates precluded full-sample time-resolved x-ray data collection. Oocytes can be illuminated using an x-ray beam comparable to their size, but then position-dependent information that allows distinguishing of surface frost and interior ice is lost. Coherent x-ray imaging techniques^27^ can yield 3D maps of a sample, but acquiring the required data during the short warming transient imposes significant constraints.

S10. Cryoprotectant-free cryopreservation of mammalian cells

Successful cryoprotectant-free cryopreservation of mammalian cells by ink-jet printing onto a cryocooled substrate has been reported^28^. Bovine and human oocytes have much larger volumes (~500 pL) than the mouse fibroblast cells (~5 pL) and drops (~40 pL) used and so maximum achievable cooling and warming rates are lower. Solutes, including those present in the cell culture media used, still suppress ice nucleation and growth, even though they may not be labelled as cryoprotectants^3,29^. High speed videos of drop deposition on glass substrates cooled from below by liquid nitrogen show drop solidification before substrate contact and a complete absence of drop deformation on impact, implying that the important cooling occurred before impact and that actual cooling rates were much smaller than the 2.2 ×10^6^ °C/min claimed based on simulations of heat transfer from a drop in full contact with a cryogenically cooled substrate. No assays are performed that robustly probe ice formation after cooling or during warming. Observed Raman peak shifts (which are not analyzed by fitting) and optical images of cold drops do not rule out the presence of a substantial ice fraction, and substantial ice likely formed during warming.

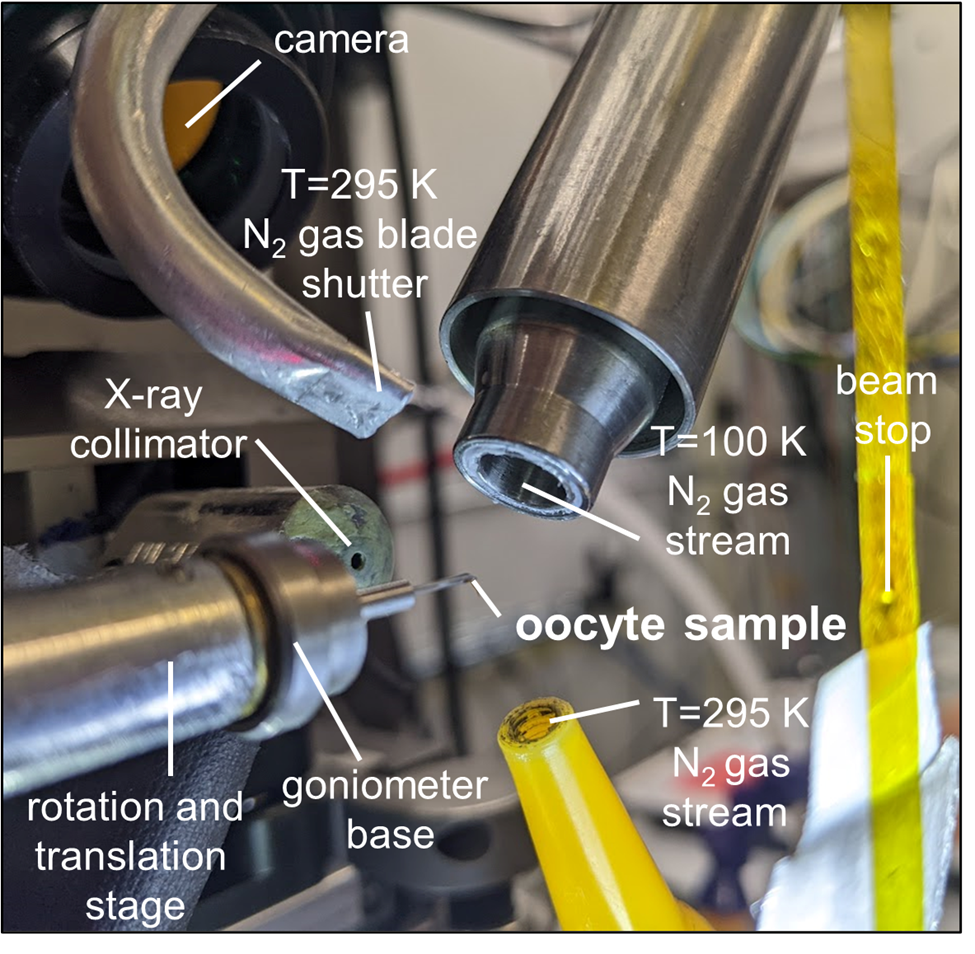

**Figure S1. Experimental setup for measuring diffraction from oocytes after cooling and during thawing at CHESS beamline 7IDB2.** Oocyte samples on crystallography supports or on Cryotop tips inserted into goniometer bases were transferred in a T<100 K automounter gripper to a rotation stage, where they were cooled by a T=100 K N_2_ gas stream. A separate room temperature N_2_ gas supply was connected through a solenoid valve to a gas blade shutter – which generated a flat gas jet to block the *T*=100 K N_2_ gas stream from reaching the sample – and to a second tube which directed a jet of room temperature N_2_ gas at the sample to warm it.

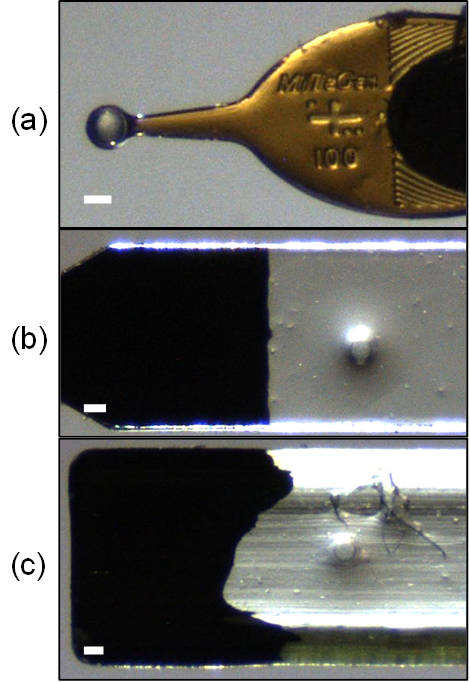

**Figure S2. Oocyte supports used for cooling and thawing.** **(a)** Crystallography support (MicroLoops LD 100, MiTeGen) with a 100 μm diameter aperture and 10 μm thick polymer in the aperture region. **(b)** Current Cryotop™ vitrification support (Kitazato), ~930 um wide with ~90 um thick polymer. **(c)** Original CryoTop™, ~1100 um wide with curved, ~200 μm thick polymer. Only the sample-accepting end of each support is shown.

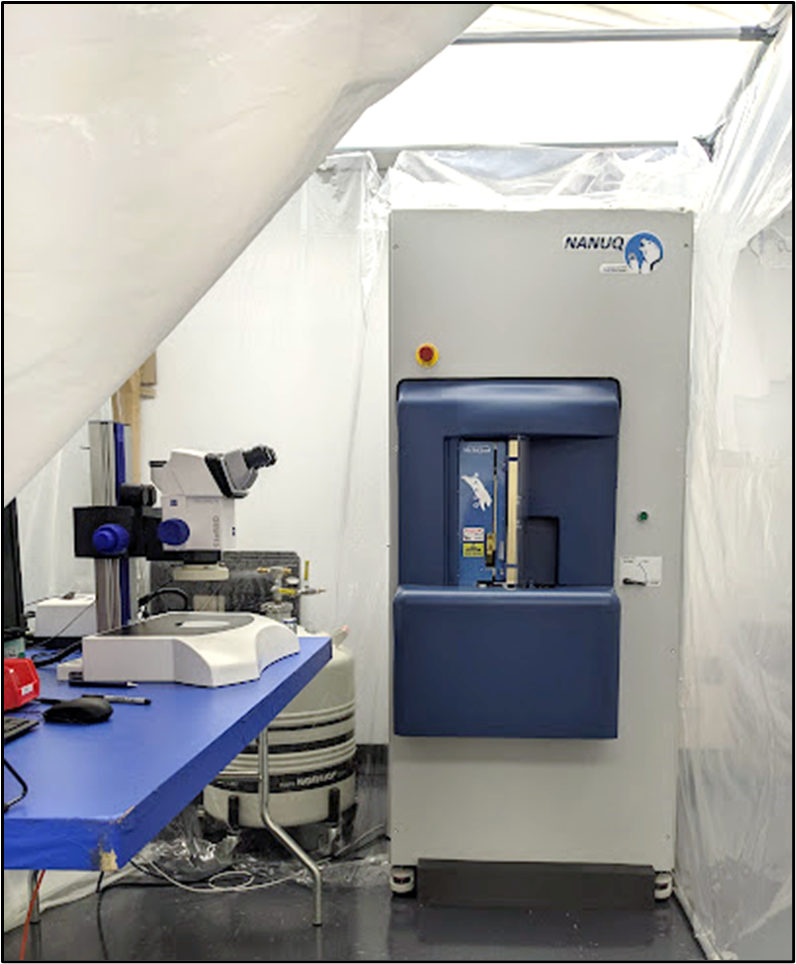

Figure S3. Experimental setup for soaking and cooling oocyte samples. An automated cryocrystallography plunge cooler (NANUQ^TM^, MiTeGen) plunged oocyte samples − on cryocrystallography supports and on Cryotop™ supports modified to insert into goniometer bases (Figure S4(b)) − at 2 m/s into liquid nitrogen at 77 K. The samples were then automatically loaded into Unipucks (Figure S4(c)). When four pucks (64 samples) had been loaded, they were removed to a separate liquid nitrogen filled tub containing a crystallography cane (Figure S4(d)). When the cane was fully loaded, it was transferred to a wide mouth Dewar filled with liquid nitrogen, or to a cryogenic dry shipping Dewar. Soaking and transfer of oocyte samples to crystallography mounts was performed as described in Section S2 adjacent to the plunge cooler and using the microscope shown. To minimize evaporation from equilibration and vitrification solutions and from oocytes during preparation, the workspace was enclosed in a “tent” humidified to ~80% r.h., and samples were immediately plunged cooled after their final soak.

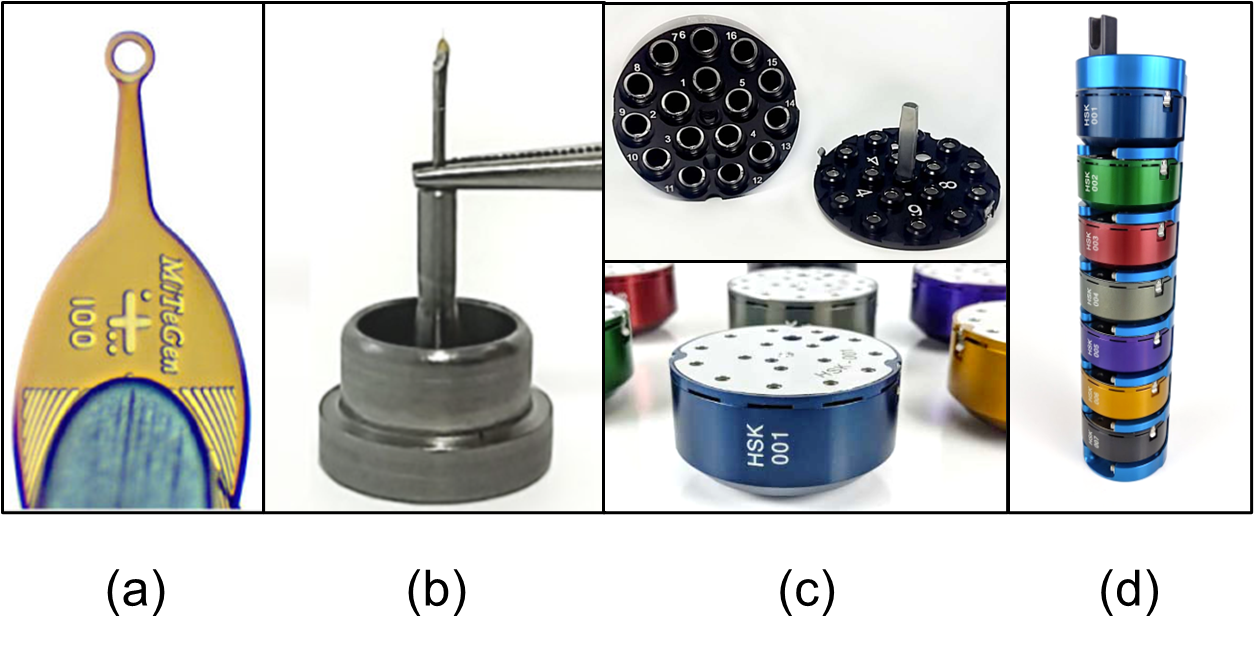

**Figure S4.** **Cryocrystallography tools and hardware adapted for use for cooling and storing oocytes.** **(a)** Crystallography support (MicroLoops LD 100, MiTeGen) with a 100 μm diameter aperture and 10 μm thick polymer in the aperture region. The polymer is attached to a beveled stainless-steel rod. **(b)** A crystallography goniometer base, machined from magnetic stainless steel, into which the crystallography support is inserted. **(c)** Cryocrystallography Unipuck in which samples on goniometer bases are stored after cryocooling. The Unipuck has magnets that capture the magnetic steel goniometer bases. **(d)** Unipuck storage cane, which holds Unipucks in standard wide-mouth liquid nitrogen Dewars or dry shippers, for storage and transport to the x-ray source.

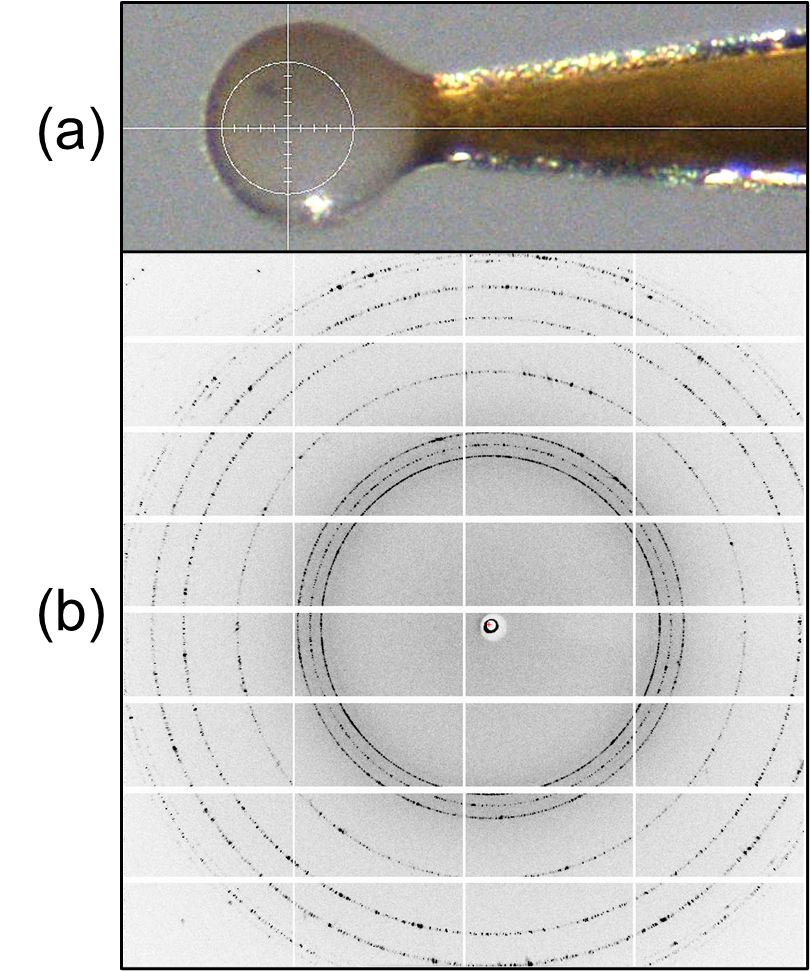

Figure S5. “Snowball” oocyte and diffraction pattern consistent with large grain hexagonal ice. The oocyte (a) was soaked in 10.5% (v/v) DMSO, 10.5% (v/v) EG, 0.35 M sucrose and “slow” cooled. At the x-ray beamline, the oocyte was initially positioned at the edge of the cryostream (by mistake), and partially thawed before it was repositioned and cooled back to *T*=100 K. (b) The cycle of slow cooling, slow thawing, and then slow cooling allowed growth of hexagonal ice crystals of much larger average size than was observed in any of our “controlled” cooling and warming experiments.

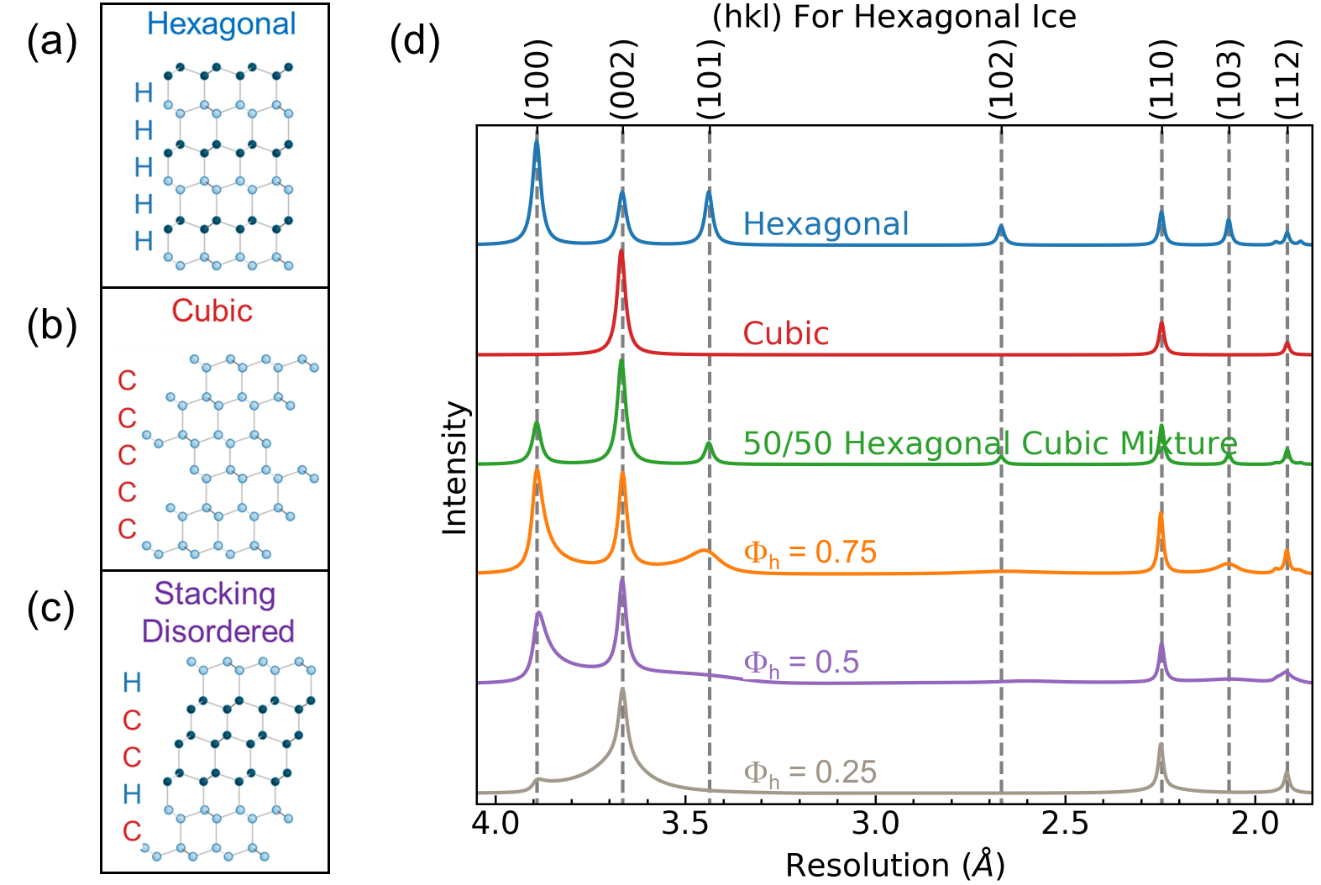

**Figure S6. Types of crystalline ice formed at ambient pressure and their corresponding x-ray diffraction patterns.** Adapted from Malkin et al.^10^ When samples are cooled slowly, hexagonal ice *I*_h_ **(a)** is obtained. When samples are cooled rapidly to cryogenic temperature, cubic ice *I*_c_ **(b)** or a mix of cubic and hexagonal ice crystallites was thought to be obtained. However, the observed ice diffraction patterns (bottom three traces in **(d)** almost always exhibit asymmetric broadening of some but not all ice diffraction peaks. Analysis and simulation show that the ice that forms during rapid cooling (under conditions of large supercooling) is in fact best modeled as a stacking disordered mix of cubic and hexagonal planes **(c)**. In the simplest model of stacking disordered ice *I*_sd_, the stacking is random, and the fraction of planes that are hexagonal is given by a parameter *Φ*_h_. As *Φ*_h_ varies from 0 (pure *I*_c_) to 1 (pure *I*_h_), the diffraction pattern evolves as shown by the lower 3 curves in **(d).**

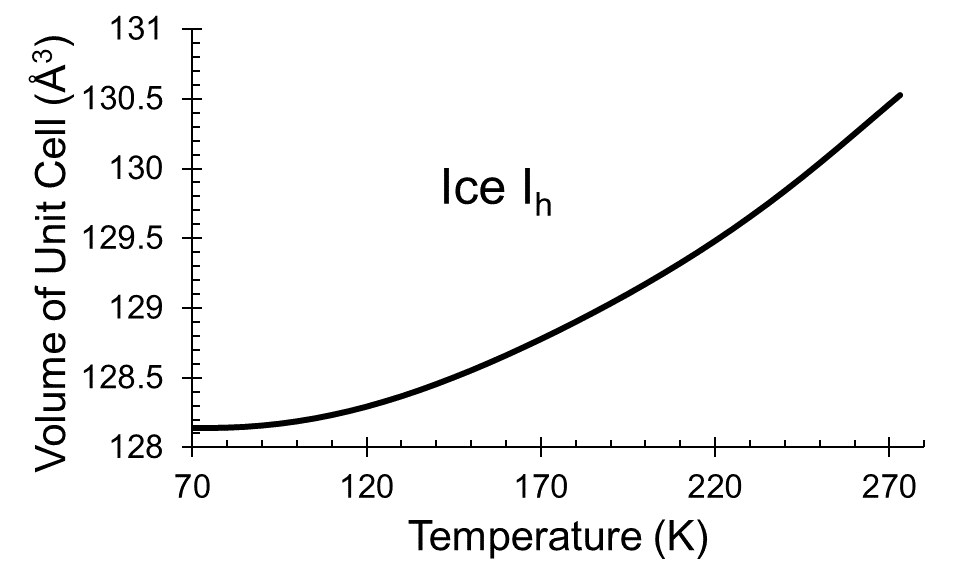

Figure S7. Unit cell volume of hexagonal ice versus temperature. The experimental volume is fit by the polynomial expression *V*_unit cell_(*T*) = A_0_ + A_1_*T* + A_2_*T*^2^ + A_3_*T*^3^ + A_4_*T*^4^ + A_5_*T*^5^ +A_6_*T*^6^ +A_7_*T*^7^, where A_0_ = 128.2147, A_1_ = 0, A_2_ = 0, A_3_ = −1.3152×10^-6^, A_4_ = 2.4837×10^-8^, A_5_ = −1.6064×10^-10^, A_6_ = 4.6097×10^-13^, and A_7_ = −4.9661×10^-16^, as determined by Rottger et al.^18,30,31^

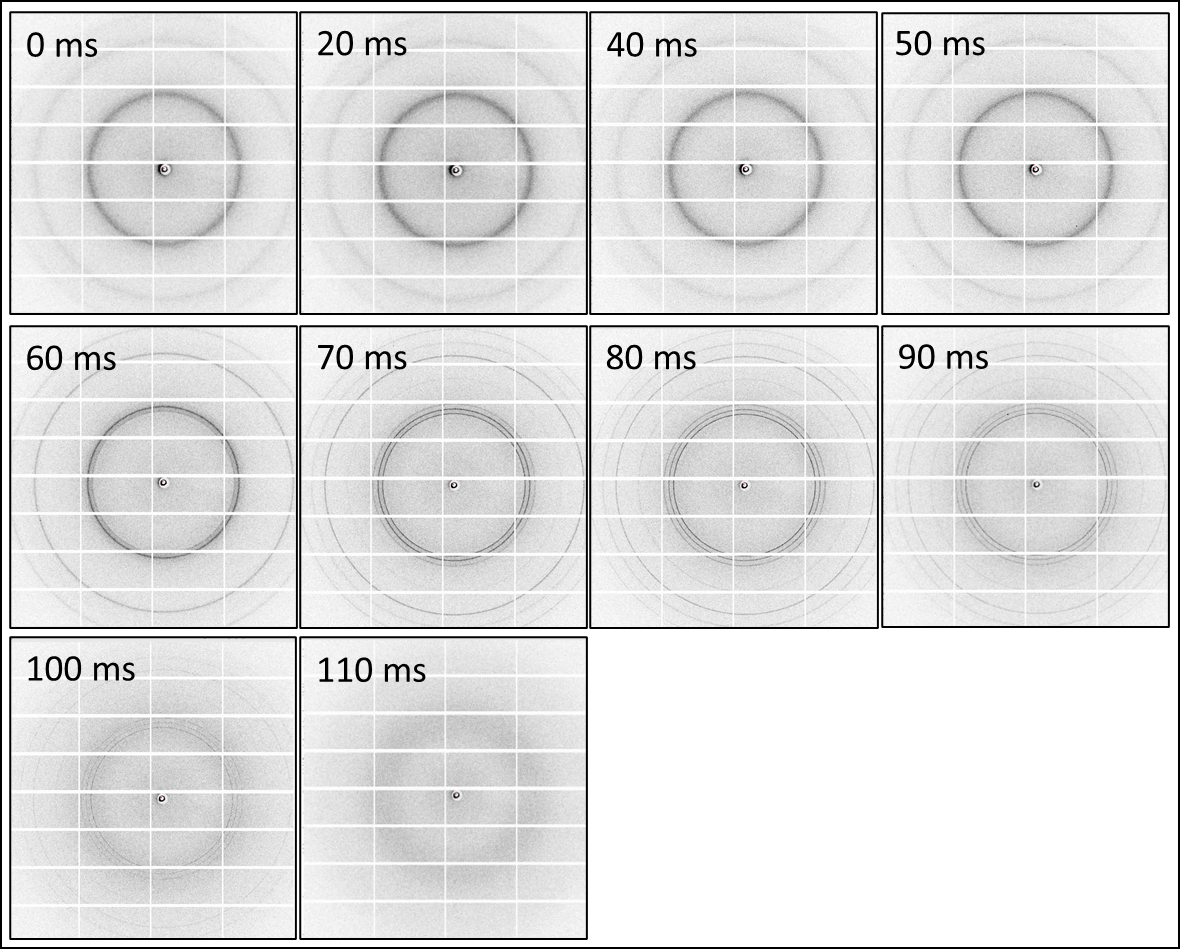

**Figure S8. Time series of diffraction images acquired during warming.** The data are for the oocyte of **Fig. 1(e),** soaked in 6% DMSO, 6% EG and 0.2 M sucrose and “fast” cooled. Diffuse rings at cubic ice resolutions evolve to sharp rings at cubic ice resolutions (60 ms) to sharp rings at all hexagonal ice positions (70-100 ms) before disappearing at 110 ms. The rings show only small azimuthal variations in intensity, suggesting that ice grain sizes are small at all times. *t*=0 corresponds to the time when solenoid valve on the room temperature N_2_ gas source, which supplied both the air blade shutter that interrupted the *T*=100 K N_2_ cryostream and the room temperature N_2_ gas stream used to warm the sample, opened. The actual start of warming was ~20-40 ms later.

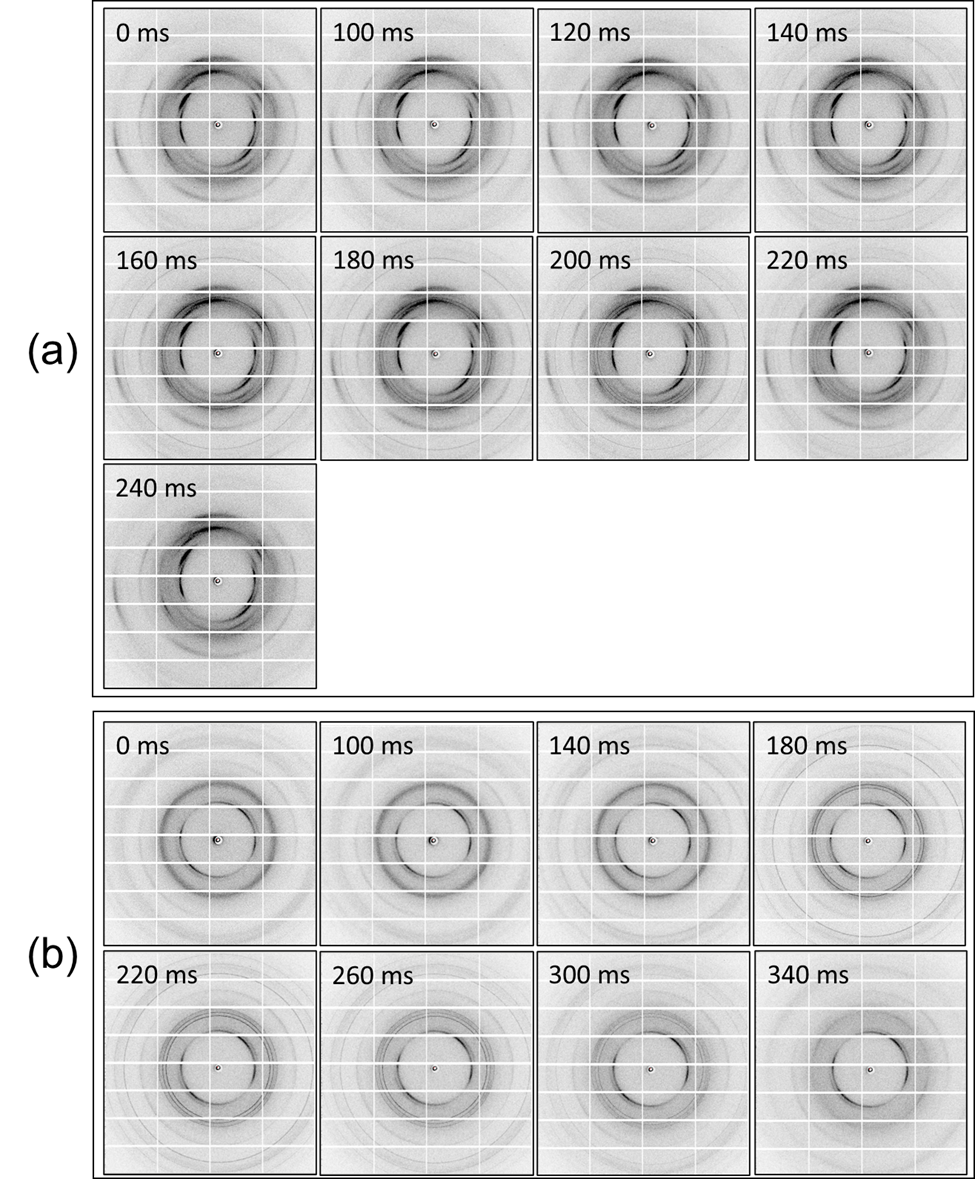

Figure S9. Time series of diffraction images acquired during warming of oocytes on thin flat Cryotops™. Unlike crystallography supports (Figure S2(a)) which have a hole beneath the central portion of the oocyte, with Cryotops (Figure S2(b)) the x-ray beam passes through the thick, highly oriented polymer, generating strongly anisotropic background scatter. (a) Oocyte soaked in 13.5% DMSO, 13.5% EG, 0.45 M sucrose and “fast” cooled. No ice diffraction was visible after cooling (0 ms). Ice rings first become visible at 140 ms and are no longer visible at 240 ms. Ice rings are largely azimuthally homogeneous, suggesting a very small grain size. (b) Oocyte soaked in 10.5% DMSO, 10.5% EG, 0. 35 M sucrose and “fast” cooled. Diffuse ice rings at cubic resolutions are visible in the cold sample at t=0. The pattern begins evolving at 140 ms, all hexagonal rings are visible at 220 ms, and ice diffraction is no longer visible at 340 ms. Significant azimuthal variation in ice ring intensity is visible at 220 ms, suggesting the presence of some larger ice grains. *t*=0 corresponds to the time when the solenoid valve on the room temperature N_2_ gas source, which supplied both the air blade shutter that interrupted the *T*=100 K N_2_ cryostream and the room temperature N_2_ gas stream used to warm the sample, opened. The actual start of warming was ~20-40 ms later.

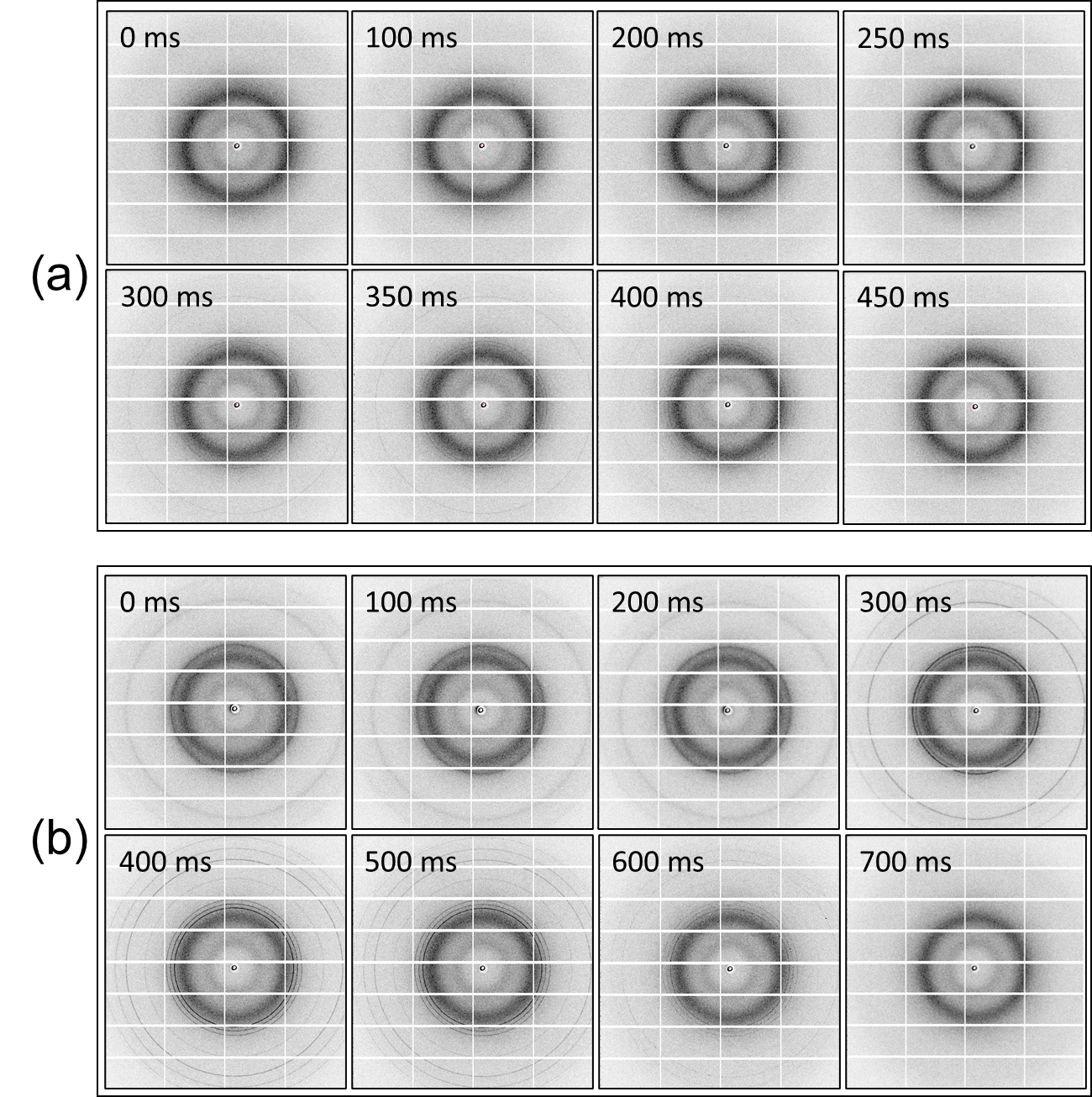

**Figure S10.** **Time series of diffraction images acquired during warming of oocytes on thicker, curved Cryotops™**. Again, the x-ray beam passes through the very thick polymer beneath the oocyte, producing very strong background diffraction. Unlike the polymer used in thin, flat Cryotops, the polymer is largely unoriented so its diffraction is azimuthally homogeneous. **(a)** Oocyte soaked in 13.5% DMSO, 13.5% EG, 0.45 M sucrose and “fast” cooled. No ice diffraction was visible after cooling (0 ms). Ice rings first become visible at 300 ms, and are no longer visible at 450 ms. Ice rings are largely azimuthally homogeneous, suggesting a very small grain size. **(b)** Oocyte soaked in 9% DMSO, 9% EG, 0.3 M sucrose and “fast” cooled. Somewhat diffuse Ice rings at cubic resolutions are visible in the cold sample at *t*=0. The pattern begins evolving at ~300 ms, all hexagonal rings are visible at 400 ms, and ice diffraction is no longer visible at 700 ms. Significant azimuthal variation in ice ring intensity is visible at *t*=300-600 ms, suggesting the presence of some larger ice grains. *t*=0 corresponds to the time when the solenoid valve on the room temperature N_2_ gas source, which supplied both the air blade shutter that interrupted the *T*=100 K N_2_ cryostream and the room temperature N_2_ gas stream used to warm the sample, opened. The actual start of warming, as indicated by the beginning of the increase in ice unit cell in samples that developed ice on cooling, was ~20-40 ms later.

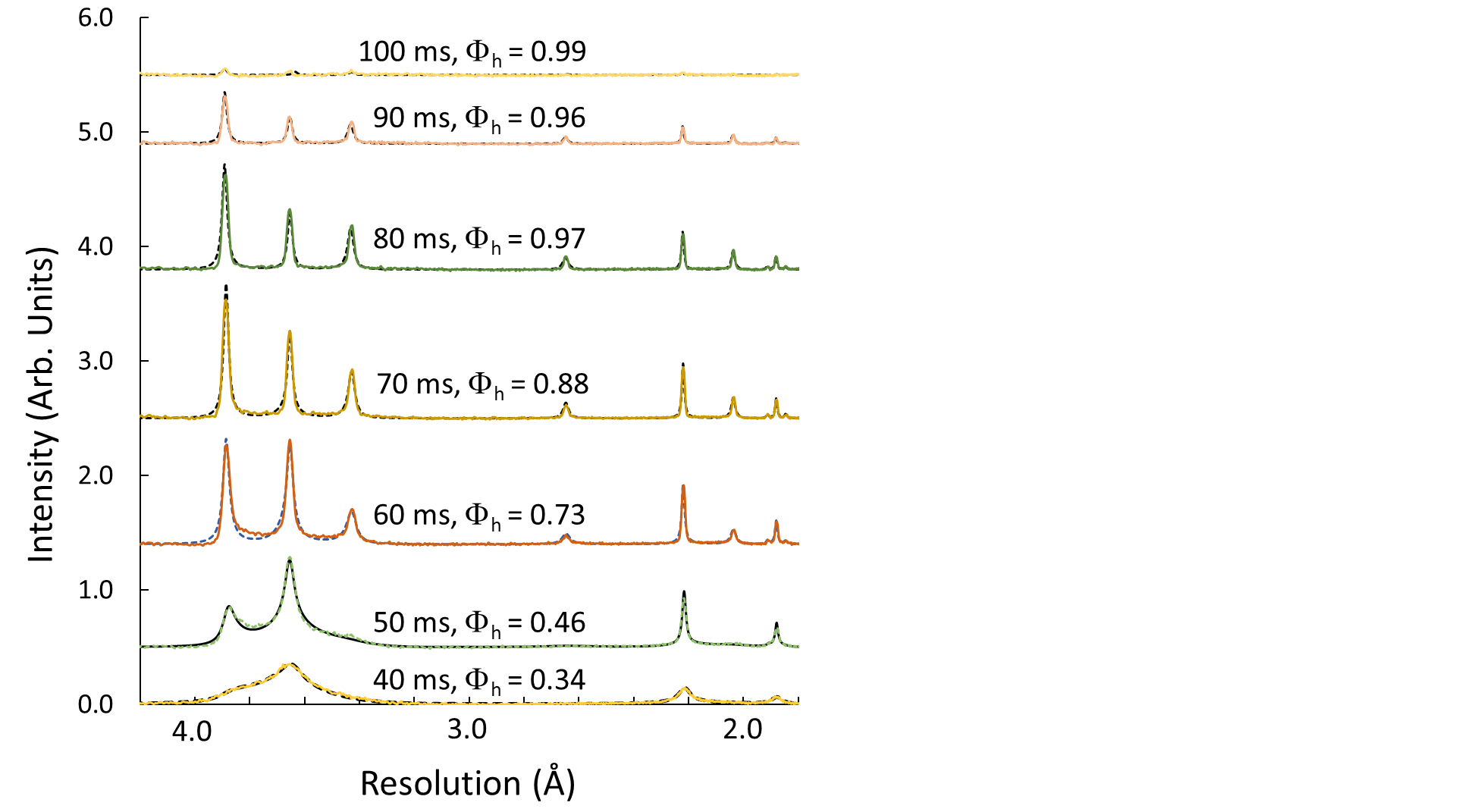

(a)

(b)

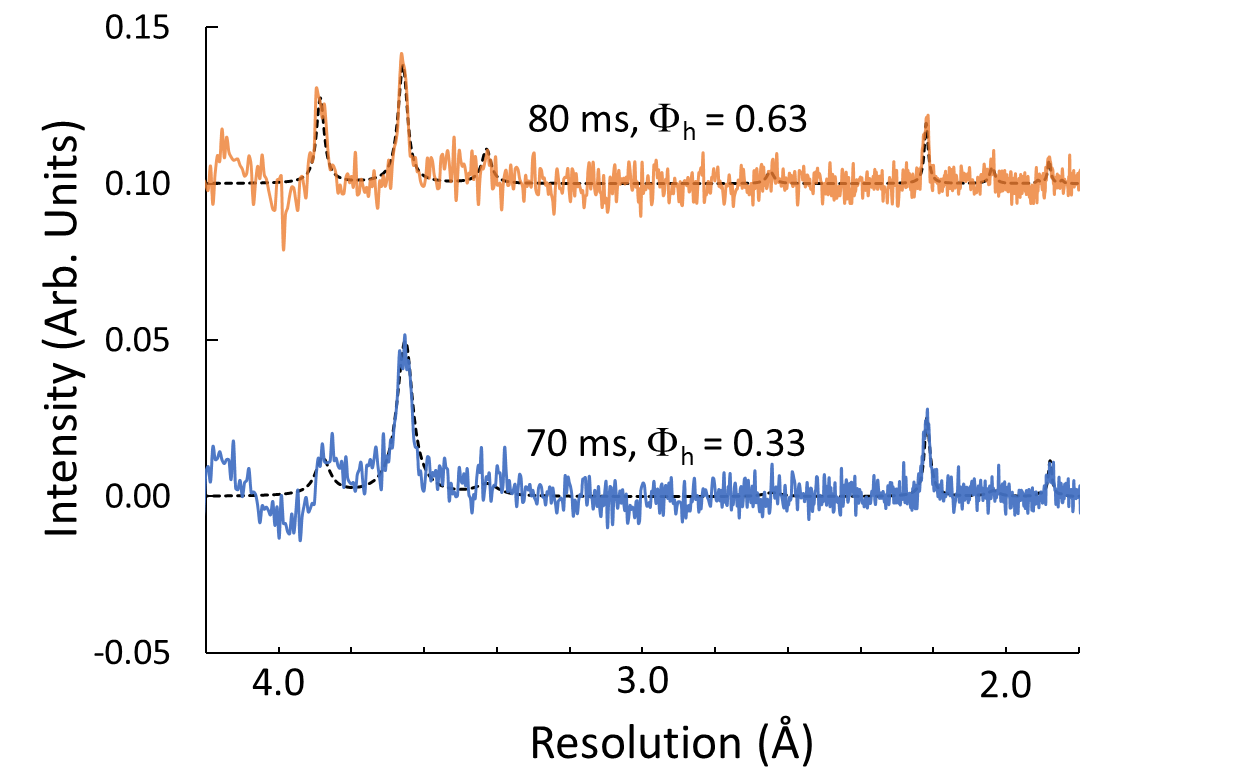

**Figure S11. Evolution of oocyte diffraction during warming.** Azimuthally integrated and background subtracted intensity for the oocyte of **(a)** **Figure 5(a)**, soaked in 7.5% DMSO, 7.5% EG, and 0.25 M sucrose, moved through oil to remove surface solvent, and “fast” cooled; and **(b)** **Figure 5(b)**, soaked in 15% DMSO, 15% EG, and 0.5 M sucrose and “fast” cooled. The black dashed lines show DIFFaX fits to a model of stacking disordered ice with fit parameters Φ_h_ shown. Note that the intensity scale in (b) is roughly 20 times smaller than in (a); the maximum ice diffraction intensity in (b) was extremely weak – ~4% of that in (a) – indicating a tiny maximum ice fraction. For a 100 μm oocyte, a ~5 μm thick layer of surface solution (vitrification solution) would occupy ~10% of the x-ray illuminated volume. The observed ice diffraction in (b) most likely arose from the surface solvent.

**Table S1. Equilibration and vitrification solution compositions used to prepare oocytes.** Results presented in the manuscript focus on the first 7 conditions.

| Condition # | Equilibration solution | | | Vitrification solution | | |
| --- | --- | --- | --- | --- | --- | --- |
|  | DMSO | EG | sucrose | DMSO | EG | sucrose |
| 1 | 7.50% | 7.50% | 0% | 15.00% | 15.00% | 0.50 M |
| 2 | 6.75% | 6.75% | 0% | 13.50% | 13.50% | 0.45 M |
| 3 | 6.00% | 6.00% | 0% | 12.00% | 12.00% | 0.40 M |
| 4 | 5.25% | 5.25% | 0% | 10.50% | 10.50% | 0.35 M |
| 5 | 4.50% | 4.50% | 0% | 9.00% | 9.00% | 0.30 M |
| 6 | 3.75% | 3.75% | 0% | 7.50% | 7.50% | 0.25 M |
| 7 | 3.00% | 3.00% | 0% | 6.00% | 6.00% | 0.20 M |
| 8 | 7.50% | 7.50% | 0% | 15.00% | 15.00% | 0.5 M |
| 9 | 6.75% | 6.75% | 0% | 13.50% | 13.50% | 0.5 M |
| 10 | 6.00% | 6.00% | 0% | 12.00% | 12.00% | 0.5 M |
| 11 | 5.25% | 5.25% | 0% | 10.50% | 10.50% | 0.5 M |
| 12 | 4.50% | 4.50% | 0% | 9.00% | 9.00% | 0.5 M |
| 13 | 3.75% | 3.75% | 0% | 7.50% | 7.50% | 0.5 M |
| 14 | 3.00% | 3.00% | 0% | 6.00% | 6.00% | 0.5 M |

**Table S2.** **Miller indices and resolutions of hexagonal ice I_h_ diffraction rings at resolutions numerically larger than 1.5 Å.** Ice rings at the resolutions indicated in shaded rows are not broadened by stacking disorder. Cubic ice I_c_ generates diffraction rings only at these resolutions; the additional peaks of hexagonal ice are broadened by stacking disorder.

| **(hkl)** | **Resolution (Å)** |
| --- | --- |
| (100) | 3.895 |
| (002) / c111 | 3.661 |
| (101) | 3.438 |
| (102) | 2.667 |
| (110) / c220 | 2.249 |
| (103) | 2.068 |
| (200) | 1.947 |
| (112) / c311 | 1.916 |
| (201) | 1.882 |
| (202) | 1.719 |
| (203) | 1.522 |
